# Fluoride export is required for competitive fitness of pathogenic microorganisms in dental biofilm models

**DOI:** 10.1101/2024.01.18.576223

**Authors:** Aditya Banerjee, Chia-Yu Kang, Minjun An, B. Ben Koff, Sham Sunder, Anuj Kumar, Livia M. A. Tenuta, Randy B. Stockbridge

## Abstract

Microorganisms resist fluoride toxicity using fluoride export proteins from one of several different molecular families. Cariogenic species *Streptococcus mutans* and *Candida albicans* extrude intracellular fluoride using a CLC^F^ F^−^/H^+^ antiporter and FEX fluoride channel, respectively, whereas commensal eubacteria, such as *Streptococcus gordonii,* export fluoride using a Fluc fluoride channel. In this work, we examine how genetic knockout of fluoride export impacts pathogen fitness in single-species and three-species dental biofilm models. For biofilms generated using *S. mutans* with genetic knockout of the CLC^F^ transporter, exposure to low fluoride concentrations decreased *S. mutans* counts, synergistically reduced the populations of *C. albicans*, increased the relative proportion of commensal *S. gordonii*, and reduced properties associated with biofilm pathogenicity, including acid production and hydroxyapatite dissolution. Biofilms prepared with *C. albicans* with genetic knockout of the FEX channel also exhibited reduced fitness in the presence of fluoride, but to a lesser degree. Imaging studies indicate that *S. mutans* is highly sensitive to fluoride, with the knockout strain undergoing complete lysis when exposed to low fluoride for a moderate amount of time, and biochemical purification the *S. mutans* CLC^F^ transporter and functional reconstitution establishes that the functional protein is a dimer encoded by a single gene. Together, these findings suggest that fluoride export by oral pathogens can be targeted by specific inhibitors to restore biofilm symbiosis in dental biofilms, and that *S. mutans* is especially susceptible to fluoride toxicity.

**Importance:** Dental caries is a globally prevalent condition that occurs when pathogenic species, including *Streptococcus mutans* and *Candida albicans*, outcompete beneficial species, such as *Streptococcus gordonii,* in the dental biofilm. Fluoride is routinely used in oral hygiene to prevent dental caries. Fluoride also has antimicrobial properties, although most microbes possess fluoride exporters to resist its toxicity. This work shows that sensitization of cariogenic species *Streptococcus mutans* and *Candida albicans* to fluoride by genetic knockout of fluoride exporters alters the microbial composition and pathogenic properties of dental biofilms. These results suggest that the development of drugs that inhibit fluoride exporters could potentiate the anticaries effect of fluoride in over-the-counter products like toothpastes and mouth rinses. This is a novel strategy to treat dental caries.

## Introduction

Dental caries is a globally prevalent disease that affects almost 2.4 billion adults and 621 million children (1, 2). In addition to causing pain and stress, infections resulting from tooth decay can cause 0.1% fatality among patients (3). Dental caries is caused by a pathogenic dental biofilm composed of acidogenic and aciduric species like *Streptococcus mutans* and *Candida albicans*, which feed on a diet rich in fermentable carbohydrates (4). *S. mutans* and *C. albicans* are symbionts, forming mixed species biofilms that are fitter and more cariogenic than single species biofilms(5–7). Under sugar exposure, these cariogenic species reduce the biological diversity of the dental biofilm and outcompete oral eubiota like the commensal streptococci *S. gordonii*, *S. sanguinis*, and *S. oralis* (8).

Fluoride is a widely accepted anticaries agent present in toothpastes and other dental healthcare products that promotes tooth remineralization (9). Fluoride also exhibits antimicrobial activity due to its broad-spectrum inhibition of essential metabolic enzymes, including enolase in the glycolytic pathway, pyrophosphatase, and ATP-consuming enzymes (10). Over-the-counter fluoride products like toothpastes and rinses contain fluoride in the range of 12 to 60 mM fluoride (226 to 1100 ppm F^−^), which is high enough to have such antimicrobial effects(11). However, during the first hour after using these treatments, fluoride concentrations in the oral fluids drop to <1 mM (12–15). Because most microbes possess fluoride export proteins to maintain cytoplasmic fluoride at low levels(16), at these concentrations the antimicrobial effect is clinically irrelevant. Genetic knockout of fluoride exporters causes fluoride hypersensitivity in multiple species, including oral microbiota, reducing the inhibitory concentration of fluoride to the tens-to-hundreds of micromolar range (17–19). This enhanced toxicity suggests that inhibition of fluoride efflux by pathogenic microbes could potentiate the effects of oral fluoride and reduce dental dysbiosis at clinically relevant fluoride concentrations and exposure times.

Two additional factors suggest that it might be feasible to specifically target fluoride efflux by oral pathogens. First, microbial fluoride sensitivity is amplified by low pH, as is characteristic of dysbiotic dental biofilms (20). This is because fluoride is a weak acid (pK_a_ of 3.4), and its conjugate acid, HF, can readily cross the cell membrane, where it dissociates into H^+^ and F^−^ in the relatively higher pH of the cytoplasm (21). In the absence of a fluoride export mechanism, these impermeable ions become trapped and accumulate intracellularly (20). Second, pathogenic and health-associated oral species possess fluoride export proteins from different molecular families (**Figure 1A**). *S. mutans* possesses two genes that encode F^−^/H^+^ antiporters from the CLC (**<u>c</u>**h**l**oride **<u>c</u>**hannel) family of anion channels and transporters, termed CLC^F^ (22). In contrast, oral eubiota including *S. gordonii, S. oralis*, and *S. sanguinis* all export F^−^ via the action of Fluc (**<u>Flu</u>**oride **<u>c</u>**hannel) proteins (also known by their gene name, *crcB*), which harness the electrical component of the proton motive force to expel cytoplasmic fluoride (23). Eukaryotes, including *C. albicans*, express a third type of fluoride exporter called FEX (**<u>F</u>**luoride **<u>Ex</u>**porter) (18). These channels are structurally related to the Flucs, but possess a more complicated two-domain fold (24, 25).

**Fig. 1.**
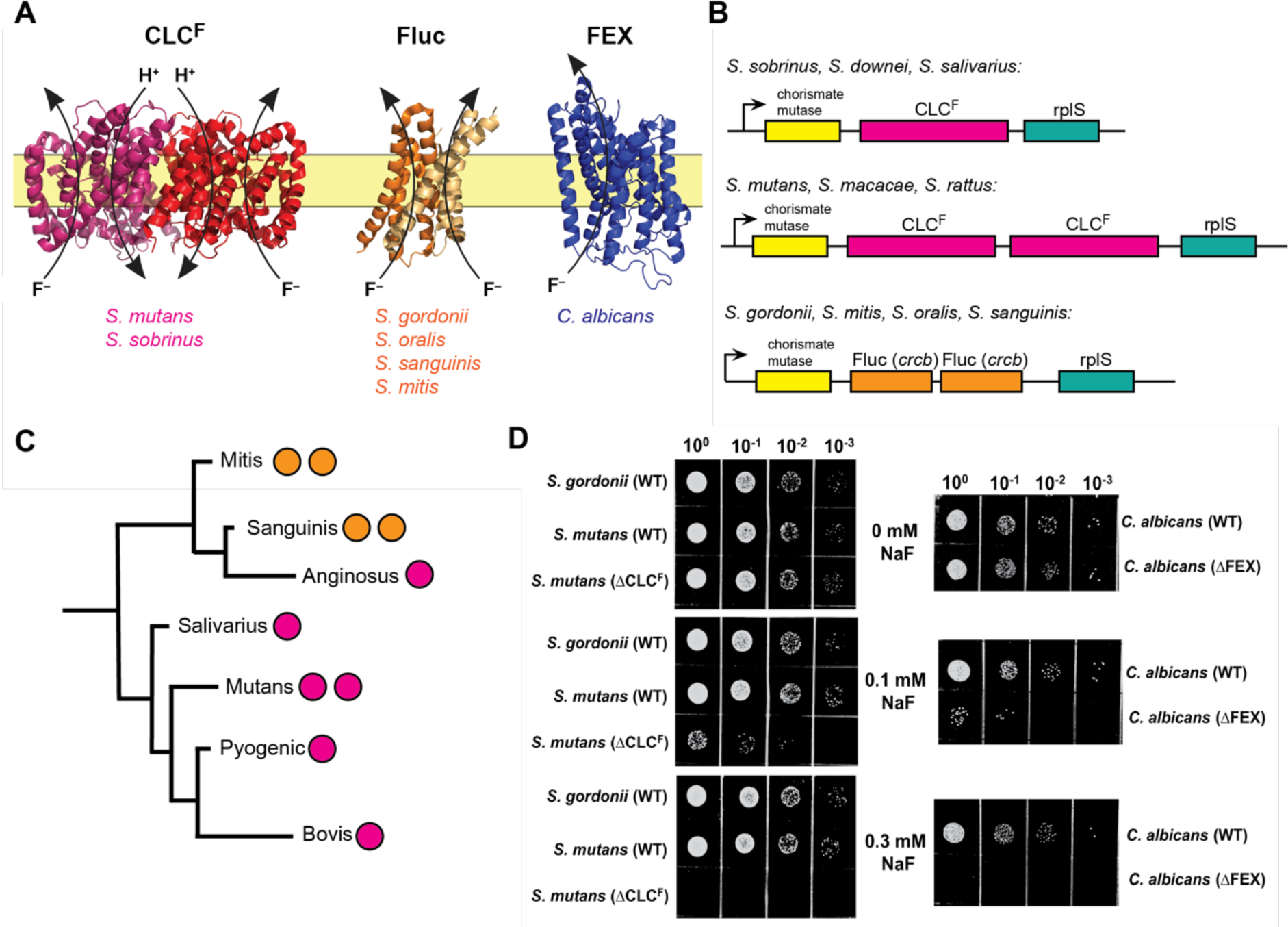
Fluoride exporters from representative oral microorganisms. **A.** Structures of fluoride exporters: F^−^/H^+^ antiporter CLC^F^ (PDB: 6D0J (49)); F^−^ channel Fluc (PDB: 5NKQ (50)); and F^−^ channel FEX (AlphaFold model). Representative oral species that possess each type of exporter are indicated. **B.** Operon architecture of selected oral streptococci. **C.** Phylogenetic tree of major streptococcus groups with the number and kind of fluoride exporter genes indicated (CLC^F^: magenta; Fluc: orange). Phylogenetic relationship and branch lengths from (51). **D.** 10-fold serial dilutions of *S. mutans*, *S. gordonii,* and *C. albicans* on plates containing NaF at the indicated concentration.

In this work, we test our hypothesis that elimination of fluoride export in cariogenic *S. mutans* and *C. albicans* reduces their competitive fitness in mixed species biofilms, with a concomitant reduction in cariogenic biofilm properties. We examine the species composition and pathogenic potential of dental biofilms composed of *S. mutans*, *C. albicans,* and the beneficial commensal *S. gordonii*. We find that genetic knockout of fluoride exporters in the pathogenic species yields biofilms with a larger proportion of commensal bacteria, lower total biomass and reduced pathogenicity at fluoride concentrations as low as 0.2 mM (3.8 ppm F^−^). We further show that *S. mutans* is especially sensitive to the ablation of fluoride efflux, as the CLC^F^ knockout strain exhibits complete lysis upon exposure to 1 mM fluoride for moderate lengths of time. Biochemical and genetic analysis of the two CLC^F^-encoding genes in *S. mutans* shows that only one of the two open reading frames contributes to fluoride export by this pathogen. Together, our data indicate that fluoride exporters are potential targets for inhibition to eliminate cariogenic bacteria, reduce dental biofilm dysbiosis, and improve oral health.

## Materials and Methods

### Microbial strains

Experiments were performed with *S. mutans* UA159 (ATCC 700610), *S. gordonii* V288 (ATCC 35105), and *C. albicans* BWP17. For *S. mutans,* the SMU.1289 and SMU.1290 open reading frames were deleted by homologous recombination. For the double knockout strain, SMU.1290 and SMU.1289 were deleted sequentially (referred to in the text as ΔCLC^F^). Synthetic genes encoding the spectinomycin and erythromycin resistance genes were prepared with sequence homology to the upstream and downstream regions of 1290 and 1289, respectively, using two-step PCR following (26). Primers are reported in **Supplementary Table 1**. Electrocompetent *S. mutans* cells were electroporated with 1 µg of the disruption construct, incubated for 2 h in TSB supplemented with 1% (w/v) glucose at 37°C under 5% CO_2_ for 2 h, and plated on tryptic soy agar containing 1% (w/v) glucose and the appropriate antibiotic. Plates were incubated at 37°C under 5% CO_2_ for 2-6 days until colonies appeared. Mutants were confirmed using PCR analysis.

For *C. albicans*, a homozygous FEX deletion (ΔΔ*FEX1*, referred to in the text by the shorthand ΔFEX) was prepared by homologous recombination-based gene replacement. Splice overlap-extension PCR was used to generate a cassette with *HIS1* from *C. dubliniensis* flanked by approximately 300 base pairs of sequence from the region immediately upstream and downstream of the targeted fluoride exporter gene(27, 28). The *HIS1* knockout cassette was introduced into *C. albicans* BWP17 (*ura3*::*imm434*/*ura3*::*imm434 iro1*/*iro1*::*imm434 his1*::*hisG*/*his1*::*hisG arg4*/*arg4*) by standard methods of lithium acetate-mediated DNA transformation(29). The site of *CdHIS1* integration in selected transformants was verified by PCR. A homozygous deletion mutant was generated by replacement of the second gene copy using a *SAT1* deletion cassette with flanking sequence homology to the targeted fluoride exporter gene. The *SAT1* knockout cassette was generated as described above. Transformants were selected for growth on yeast synthetic dropout medium without histidine and with 100 µg/mL nourseothricin. Cassette integration was again verified by PCR, and retention of the *HIS1* cassette was also verified in this transformant. The homozygous deletion mutant was subsequently archived as a 15% glycerol stock at −80°C.

### Media and growth conditions

Typically, *S. mutans* was cultured in tryptic soy broth (TSB) with 1% glucose (w/v) at 5% CO_2_ and 37°C. For Δ1289 and Δ1290 *S. mutans*, media was supplemented with 10 µg/mL erythromycin or 300 µg/mL spectinomycin, respectively, and with both antibiotics for the Δ1290Δ1289 double knockout strain (ΔCLC^F^). *C. albicans* was typically cultured in yeast peptone dextrose (YPD) (Sigma-Aldrich, St. Louis, MO) supplemented with 25 µg/mL uridine aerobically at 30°C.

For dilution assays, overnight cultures were diluted to adjust the OD_600_ to 0.1. Ten-fold serial dilutions were spotted on tryptic soy agar with 1% (w/v) glucose (for *S. mutans*) or Sabouraud dextrose agar (for *C. albicans*) containing NaF as indicated in the text. For the *S. mutans* fluoride survival assays, fresh cultures of ΔCLC^F^ or WT *S. mutans* were grown in TSB with 1% (w/v) glucose with antibiotic as appropriate to OD_600_ of 0.3 at 37°C and 5% CO_2_. 50 µL of the culture was used to inoculate 5 mL of fresh media containing 0.3 mM or 1 mM NaF. Cultures were anaerobically incubated at 37°C for the indicated time, at which point 5 µL of the culture was collected and immediately spread onto tryptic soy agar containing 1% (w/v) glucose without any NaF. CFUs were counted after 48h incubation.

### Transmission electron microscopy

ΔCLC^F^ and WT *S. mutans* were incubated for 4 h in TSB supplemented with 1% (w/v) glucose and NaF. After pelleting, cells were fixed overnight at 4°C in glutaraldehyde (2% (v/v) in 0.1 M Sorensen’s buffer, pH 7.4). Cells were washed multiple times in 0.1 M Sorensen’s buffer and then post-fixed in 1% (w/v) osmium tetroxide for 2h at room temperature. After additional washes with 0.1 M Sorensen’s buffer, cells were immobilized in 1% (w/v) agarose. Agarose slices (∼0.5 mm) were dehydrated using four 15-min washes with stepwise increases in ethanol from 25%-100%, and washed three times in 100% propylene oxide. Embedding resin was prepared by mixing 30g Poly/Bed^®^812 (PolySciences Inc., USA), dodecenylsuccinic anhydride (DDSA), and nadic methyl annhydride (NMA) in a 2:1:1 ratio in propylene oxide. 2% Tris(dimethylaminomethyl)phenol (DMP-30) was added to the resin as an accelerator. Dehydrated agarose slices were infiltrated with resin in three concentration steps (25%, 50%, and 75%), with 16h room temperature incubation for each. A final incubation was performed at 60°C under vacuum for 48h in electron microscopy molds. After polymerization, 50 nm sections were obtained using a Leica UC7 ultramicrotome, loaded onto copper grids, stained in 7% (w/v) uranyl acetate for 10 min, washed in deionized water for 10 min and post-stained in Reynold’s lead citrate for 5 min. Imaging was performed on a JEOL 1400-plus transmission electron microscope equipped with an XR401 AMT sCMOS camera.

### Three-species surface biofilm

Polystyrene stalks (0.25 inch diameter and 0.5 inch in length; United States Plastic Corp, USA) were attached to the inner surface of the lid of a 24-well microtiter plate using double sided tape and sterilized by ethylene oxide. Inoculum (2×10^8^, 2×10^8^ and 2×10^6^ CFU/mL for *S. mutans*, *S. gordonii* and *C. albicans,* respectively (8) in TSB supplemented with 1% (w/v) sucrose) was incubated anaerobically at 5% CO_2_ and 37°C for 8 h for initial adhesion to the polystyrene stalks. Stalks were rinsed three times for 10s in saline (0.9% (w/v) NaCl) prior to initiating the experiment, which typically used TSB supplemented with 1% (w/v) sucrose, and NaF or NaCl. Biofilms were grown for 16h at 37°C under 5% CO_2_ without antibiotics. To harvest biofilms, the stalks were rinsed in saline, detached from the lid, and immersed in sterile saline solution in microcentrifuge tubes for sonication using a cup horn sonicator.

Serial dilutions of the suspension were plated on Sabouraud dextrose agar (for *C. albicans;* aerobic 30° C incubation, 48 h), Mitis Salivarius agar (for *S. mutans*), or TSB with 1% (w/v) glucose (for *S. mutans* and *S. gordonii,* at 5% CO_2_, 37°C incubation, 24 h). To determine biofilm dry weight, the biofilm suspension was pelleted and dried at 37°C for 3 days.

### Biofilm growth on hydroxyapatite (HA) discs

HA discs, each 0.5 inch diameter and 0.08 inch width (Clarkson Chromatography Products Inc, USA) were affixed in a vertical position to the lid of a 24-well microtiter plate using double sided tape and sterilized using ethylene oxide as described(30). Biofilms were formed and adhered as in the polystyrene stalk experiments. After initial adhesion, the discs were washed and immersed in fresh TSB containing 0.1 mM glucose without antibiotic, supplemented with 0.2 mM NaF or NaCl and incubated at 37°C under 5% CO_2_ for 16h (famine period). The next day, the discs were rinsed in saline solution and transferred to TSB supplemented with 1% (w/v) sucrose (56 mM) supplemented with 0.2 mM NaF or NaCl and incubated for 8h (feast period). After five days of feast-and-famine cycles, the HA discs were rinsed, detached from the lid and sonicated to detach the biofilm. The suspension was serially diluted onto selective media described above. The pH of the media was assessed after each media exchange using a calibrated pH electrode. Media calcium content was determined using the colorimetric Arsenazo III reagent (31). Absorbance (650 nm) was measured using a microplate reader (SpectraMax iD3, Molecular Devices, San Jose, CA) and calcium concentration determined from a standard curve. For fluoride determination, 0.1 mL of 0.5 N HCl was added per 10 mg biofilm wet weight and extracted for 3 h with agitation. Debris were pelleted, and the supernatant neutralized with TISAB II buffer containing 0.5 N NaOH. Fluoride was measured using a fluoride electrode (96-09, Thermo Fisher Scientific, Waltham, MA) and the concentration determined from a standard curve prepared using the same reagents as the samples.

### Quantitative PCR

After initial adhesion of the biofilm (WT strains of *S. mutans*, *S. gordonii* and *C. albicans*), polystyrene stalks were rinsed and re-incubated in 2 mL of fresh TSB containing 1% (w/v) sucrose and 0.2 mM NaF or NaCl. Biofilm was harvested after 2, 4, 16 or 24h of incubation and washed three times in saline with sonication to remove EPS(32). Total RNA was isolated from the biofilm using TRIzol Max Bacterial RNA isolation kit (Thermofisher Scientific, USA) following the manufacturer’s protocol. RNA (5 µg) was treated with DNaseI (1U/µL) to remove genomic DNA contamination, then reverse transcribed using Maxima First Strand cDNA synthesis kit (Thermo Fisher Scientific) following the manufacturer’s protocol. For qPCR, gene-specific primers (**Supplementary Table 1**) and 10 ng of diluted cDNA template were used. The reaction was performed with Power SYBR Green PCR Master Mix (Thermo Fisher Scientific) in qPCR Step-One Plus detection system (Applied Biosystems, Waltham, MA) using comparative C_T_ method with endogenous control *gyrase A* used to normalize the expression variance of target genes among samples using the 2^−ΔΔC^_T_ formula (33).

### Protein purification and proteoliposome preparation

For overexpression and biochemical assays, the coding sequences of CLC^F^ ORF 1289 or ORF 1290 were separately cloned into a pASK vector with a C-terminal hexahistidine tag(22) and transformed into *Escherichia coli* (BL21-DE3). Protein expression was induced at OD_600_ 0.5 with 0.5 mg/mL anhydrotetracycline for 3h. After cell disruption by sonication, protein was extracted with 2% (w/v) decyl-β-D-maltopyranoside (DM; Anatrace, Maumee, OH) for 2 h at room temperature. The lysate was centrifuged to pellet cell debris, and the protein was purified using cobalt affinity resin (1mL/L culture, Takara Bio USA, San Jose, CA). The protein was washed with 100 mM NaCl, 20 mM imidazole, 20 mM Tris-HCl pH 8.0, 5 mM DM, and eluted in the same buffer with 400 mM imidazole, before a final size exclusion purification step (Superdex 200) in 100 mM NaCl, 10 mM NaF, 20 mM 4-(2-hydroxyethyl)piperazine-1-ethanesulfonic acid (HEPES) pH 7.5, 4 mM DM. To prepare proteoliposomes, 25 μg purified protein was mixed with 5 mg of *E. coli* polar lipid extract (Avanti Polar Lipids, Alabaster, AL) solubilized in 35 mM 3-((3-cholamidopropyl)dimethylammonio)-1-propanesulfonate (CHAPS) and dialyzed against 0.3 M KF, 15 mM HEPES pH 7.0 at room temperature for 36 h with three buffer changes. Liposomes were stored in aliquots at −80°C until the day of use.

### Protein crosslinking

0.2 mg/mL purified protein was incubated with 0.125% glutaraldehyde (∼500-fold molar excess) at room temperature. The reaction was quenched with 0.15 M Tris-HCl, pH 7.5 (10-fold molar excess) prior to analysis by SDS-PAGE.

### Fluoride efflux from proteoliposomes

After three freeze/thaw cycles, the multilamellar vesicles were extruded 21 times through a 400 nm membrane filter to form proteoliposomes. Liposomes were passed through a 1.5 mL Sephadex G20 resin column equilibrated with 0.3 M potassium isethionate, 1 mM KF, 15 mM HEPES pH 7.0 and diluted 20-fold in assay buffer in a stirred chamber. Fluoride was monitored with a fluoride electrode (Cole Palmer, USA) attached to a pH meter and digitized at a sampling frequency of 5 Hz. Transport was initiated by addition of 1 μM of the potassium ionophore valinomycin to relieve the electrical potential and permit F^−^ and K^+^ to flow down their chemical gradients. At the end of the experiment, 30 mM octyl-β-D-glucopyranoside was added to release remaining encapsulated fluoride.

### Statistical analyses

Every experiment was performed as three independent biological replicates (*n* = 3). Two-way analysis of variance (ANOVA) using ‘species’ and ‘fluoride’ as the two factors, followed by Fisher’s LSD test was performed in GraphPad Prism 10 to calculate standard error (SE) and statistical significance of the independent conditions.

## Results

### Survey of fluoride export genes among the oral streptococci

We first examined the genetic context of fluoride export proteins found among streptococcus species (**Figure 1B, C**). Primary colonizers of the oral cavity from the phylogenetically related mitis and sanguinis groups, which are usually associated with oral health, possess paired Fluc genes, which typically encode heterodimeric fluoride ion channels(34). In contrast, other oral streptococci, including pathogens *S. mutans* and *S. sobrinus* and other species from the mutans, anginosus, salivarius, and downei groups, instead possess CLC^F^-type F^−^/H^+^ antiporters (**Table 1**). Species in the mutans group feature two adjacent CLC^F^ open reading frames (1289 and 1290 in *S. mutans*) that encode proteins with ∼60% sequence identity. The genetic context of the fluoride exporters is the same for both Fluc and CLC^F^ exporters and across streptococci, with the fluoride exporter gene(s) flanked by open reading frames for chorismate mutase on the 5’ end, and by ribosomal protein rplS on the 3’ end in most species.

**Table 1.** Fluoride exporters of the oral streptococci.

| Group | Example species | Fluoride exporter | Refseq Protein ID |
| --- | --- | --- | --- |
| Mitis | *Streptococcus mitis* | Fluc heterodimer | WP_049500397.1, WP_000712114.1 |
|  | *Streptococcus oralis* | Fluc heterodimer | WP_084881387.1, WP_084881388.1 |
| Sanguinis | *Streptococcus sanguinis* | Fluc heterodimer | WP_002933579.1, WP_002933586.1 |
|  | *Streptococcus gordonii* | Fluc heterodimer | WP_061603755.1, WP_061603756.1 |
| Anginosus | *Streptococcus anginosus* | CLC<sup>F</sup> (single) | WP_268730273.1 (lacks adjacent rplS) |
| Mutans | *Streptococcus mutans* UA159 | CLC<sup>F</sup> (paired) | WP_002263156.1, WP_002263157.1 |
| Salivarius | *Streptococcus salivarius* | CLC<sup>F</sup> (single) | WP_014633220.1 |
|  | *Streptococcus sobrinus* | CLC<sup>F</sup> (single) | WP_002962072.1 |
| Downei | *Streptococcus downei* | CLC<sup>F</sup> (single) | WP_002997240.1 |

**Table 2:**
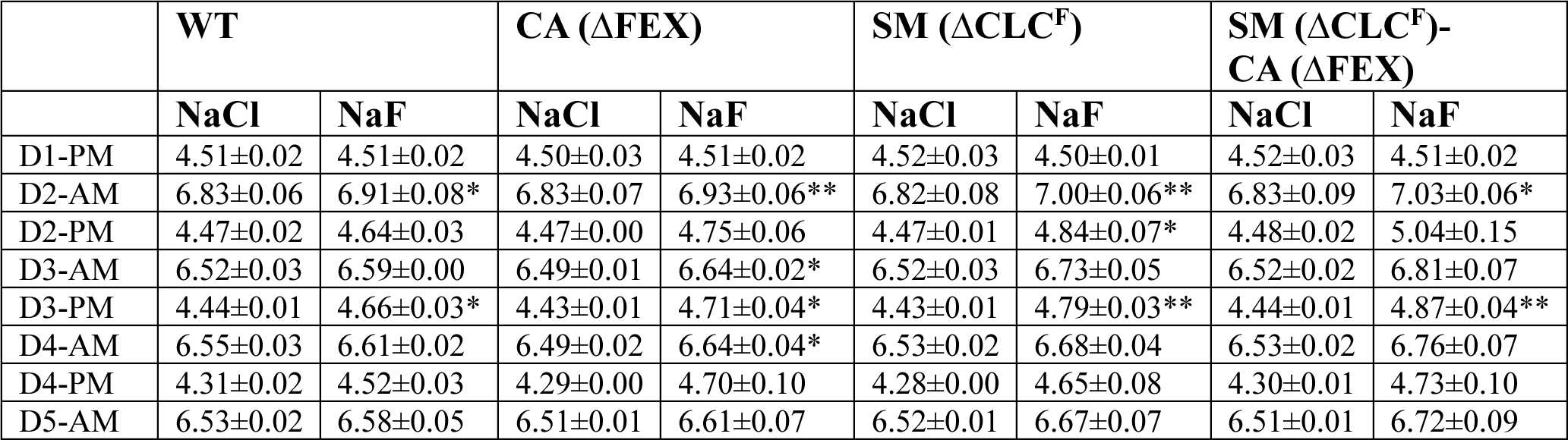
Variations in pH in the feast-and-famine model. For all data, datapoints represent mean and SEM of three independent biological replicates (*n*=3). Significance was calculated using two-way analysis of variance (ANOVA) followed by Fisher’s LSD test implemented in GraphPad Prism 8. Statistical significance is represented as ‘*’ (*p*< 0.05), ‘**’ (*p*< 0.01) and ‘***’ (*p*< 0.001).

| | WT | | CA (ΔFEX) | | SM (ΔCLC<sup>F</sup>) | | SM (ΔCLC<sup>F</sup>)-<br>CA (ΔFEX) | |
| --- | --- | --- | --- | --- | --- | --- | --- | --- |
|  | NaCl | NaF | NaCl | NaF | NaCl | NaF | NaCl | NaF |
| D1-PM | 4.51±0.02 | 4.51±0.02 | 4.50±0.03 | 4.51±0.02 | 4.52±0.03 | 4.50±0.01 | 4.52±0.03 | 4.51±0.02 |
| D2-AM | 6.83±0.06 | 6.91±0.08* | 6.83±0.07 | 6.93±0.06** | 6.82±0.08 | 7.00±0.06** | 6.83±0.09 | 7.03±0.06* |
| D2-PM | 4.47±0.02 | 4.64±0.03 | 4.47±0.00 | 4.75±0.06 | 4.47±0.01 | 4.84±0.07* | 4.48±0.02 | 5.04±0.15 |
| D3-AM | 6.52±0.03 | 6.59±0.00 | 6.49±0.01 | 6.64±0.02* | 6.52±0.03 | 6.73±0.05 | 6.52±0.02 | 6.81±0.07 |
| D3-PM | 4.44±0.01 | 4.66±0.03* | 4.43±0.01 | 4.71±0.04* | 4.43±0.01 | 4.79±0.03** | 4.44±0.01 | 4.87±0.04** |
| D4-AM | 6.55±0.03 | 6.61±0.02 | 6.49±0.02 | 6.64±0.04* | 6.53±0.02 | 6.68±0.04 | 6.53±0.02 | 6.76±0.07 |
| D4-PM | 4.31±0.02 | 4.52±0.03 | 4.29±0.00 | 4.70±0.10 | 4.28±0.00 | 4.65±0.08 | 4.30±0.01 | 4.73±0.10 |
| D5-AM | 6.53±0.02 | 6.58±0.05 | 6.51±0.01 | 6.61±0.07 | 6.52±0.01 | 6.67±0.07 | 6.51±0.01 | 6.72±0.09 |

### Fluoride is bactericidal for ΔCLC^F^ S. mutans

In agreement with the previous literature, fluoride exporter deletion strains for *S. mutans* (deletion of both the 1289 and 1290 open reading frames, referred to as ΔCLC^F^) and for *C. albicans* (homozygous FEX deletion, referred to as ΔFEX) exhibit sensitivity to fluoride. In our experiments, *C. albicans* and *S. mutans* both exhibit reduced growth by two log units in the presence of 0.1 mM NaF, and fail to grow at all in the presence of 0.3 mM NaF. The WT strains, as well as WT *S. gordonii*, are unaffected at these fluoride concentrations (**Figure 1D**). Whereas these experiments, and those of previous studies(18, 19), monitored growth of isolated strains on rich media, in the physiological context, microbes exist in complex biofilm communities. Biofilms can confer protection against environmental stressors, but are also the site of extensive interspecies competition. To evaluate fluoride sensitivity of the pathogenic strains in this context, we grew the ΔCLC^F^ *S. mutans* and ΔFEX *C. albicans* strains in three-species surface biofilms together with commensal *S. gordonii*, under conditions of increasing fluoride (**Figure 2A**). After 24 h, biofilms were harvested and replated on fluoride-free recovery plates to determine the viable CFUs of each species after fluoride treatment. Similar to the previous dilution experiment, increasing doses of fluoride between 0.1 and 0.3 mM (5.7 ppm F^−^) significantly reduced ΔFEX *C. albicans* and ΔCLC^F^ *S. mutans* CFUs harvested from the mature biofilm, whereas the number of WT *S. gordonii* CFUs remained unaffected. Exposure to 0.2 mM (3.8 ppm F^−^) fluoride reduced ΔCLC^F^ *S. mutans* and ΔFEX *C. albicans* colony counts by 830- and 50-fold respectively, compared to biofilms grown without fluoride treatment (**Figure 2A**). For ΔFEX *C. albicans*, increasing fluoride to 0.3 mM (12.6 ppm) caused an additional 4.3-fold decrease in CFUs compared with those recovered from the biofilm exposed to 0.2 mM NaF (a 200-fold decrease compared to biofilms without fluoride). This agrees with the conclusion of a previous study that, although fluoride treatment interrupts *C. albicans* growth, a fraction of the population remains viable and is able to resume growth after the fluoride challenge is removed (18). In striking contrast, we observed no viable ΔCLC^F^ *S. mutans* colonies from biofilms treated with 0.3 mM fluoride for 24h. These results suggested that ΔCLC^F^ *S. mutans* is at a severe competitive disadvantage, and perhaps killed, by relatively low concentrations of fluoride.

**Fig. 2.**
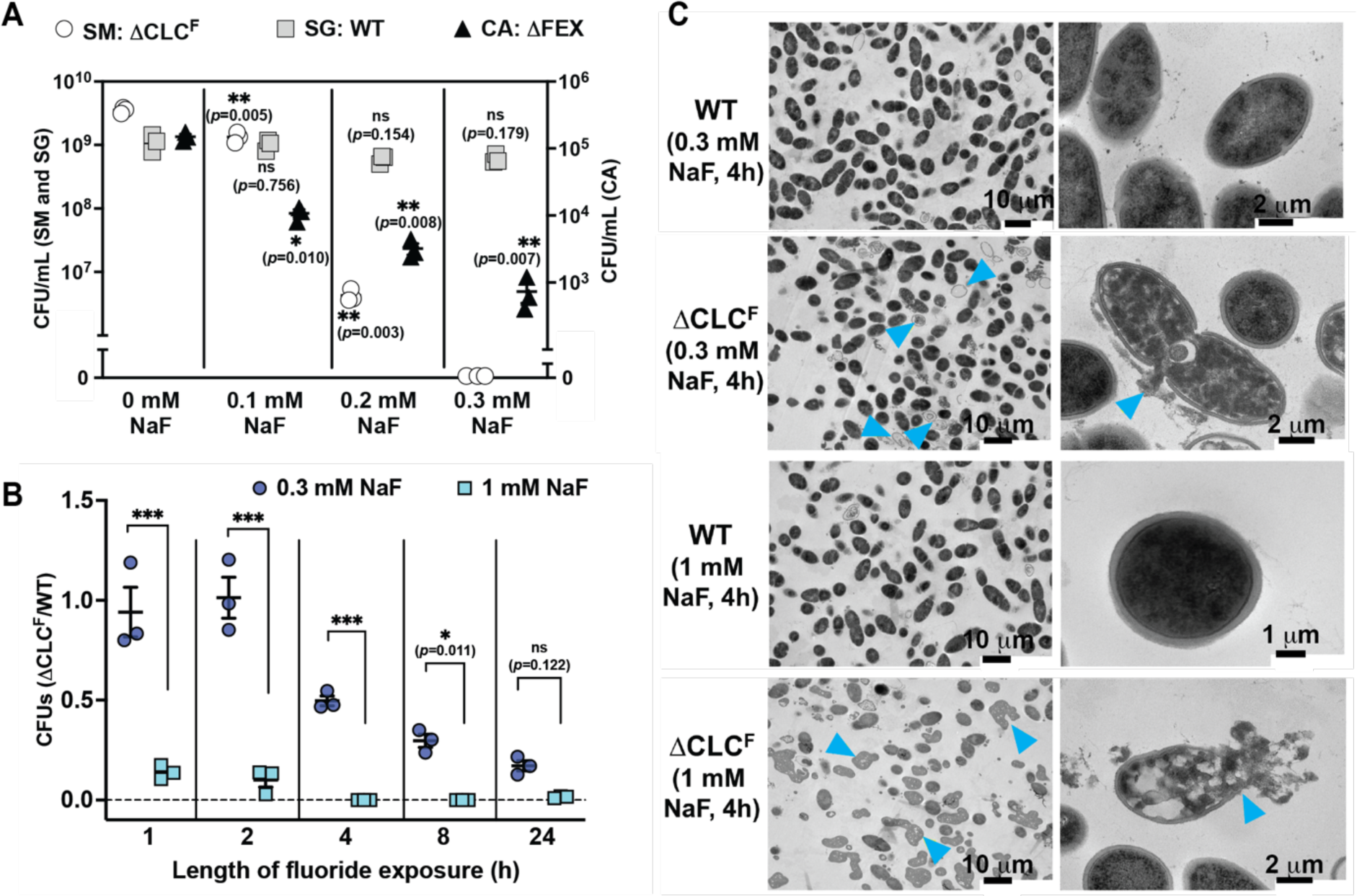
Effect of fluoride on the viability of ΔCLC^F^ *S. mutans* and ΔFEX *C. albicans*. **A.** CFUs harvested from 3-species surface biofilms (WT *S. gordonii*, ΔCLC^F^ *S. mutans*, ΔFEX *C. albicans*) after 24 h. Fluoride concentration (0, 0.1, 0.2 or 0.3 mM NaF) indicated. For biofilms grown in the presence of 0.1-0.3 mM NaF, significance is shown relative to the 0 NaF condition. Datapoints represent mean and SEM of three independent experiments (*n*=3). Significance was calculated using two-way analysis of variance (ANOVA) followed by Fisher’s LSD test. **B.** Recovered ΔCLC^F^ *S. mutans* CFUs as a function of time for samples treated with 0.3 or 1 mM NaF, expressed as a fraction relative to WT *S. mutans*. WT *S. mutans* CFUs are shown in **Supplementary Figure 1**. Datapoints represent mean and SEM of three independent experiments (*n*=3). Significance was calculated using two-way analysis of variance (ANOVA) followed by Fisher’s LSD test. ‘***’ represents *p*< 0.001. **C**. TEM images of ΔCLC^F^ or WT *S. mutans* cells after exposure to 0.3 or 1 mM NaF for 4h. Blue arrows indicate examples of ghost cells due to leak of cytoplasmic contents (0.3 mM NaF, zoomed out view), dispersed cytosolic-nucleoid material due to complete cell wall digestion (1 mM NaF, zoomed out view), or ruptures in the cell wall (zoomed in views).

To explicitly test whether fluoride treatment is bactericidal for ΔCLC^F^ *S. mutans*, we exposed planktonic cultures of WT and ΔCLC^F^ *S. mutans* to 0.3 or 1 mM NaF for increasing lengths of time before plating on fluoride-free recovery plates. For cells treated with 0.3 mM NaF, the number of viable ΔCLC^F^ *S. mutans* colonies decreased as a function of time between 4 and 24 h. Recovery of the WT strain was unaffected over this timecourse (**Supplementary Figure 1**). This effect was even more striking when cultures were treated with 1 mM NaF. After just one hour exposure, viable ΔCLC^F^ *S. mutans* CFUs were reduced 6.8-fold relative to WT (**Figure 2B**). No survival of ΔCLC^F^ *S. mutans* was detected at timepoints ≥4 h of treatment. In contrast, WT CFUs actually increased up to 24 h, indicating continued cell division (**Supplementary Figure 1**). Transmission electron microscopy (TEM) imaging showed widespread cell lysis and cell wall damage for ΔCLC^F^ *S. mutans* exposed to 0.3 mM NaF for 4 h, and lysis/total cell wall degradation of essentially all ΔCLC^F^ cells exposed to 1 mM NaF for 4 h (**Figure 2C**). WT cells exhibited normal morphology after both treatments. Together these experiments show that, if unable to export fluoride, *S. mutans* is killed and eliminated from biofilms by fluoride in the hundreds of micromolar range. The population of ΔFEX *C. albicans* is reduced by ∼two orders of magnitude by fluoride treatment in this range, but not eliminated.

### Fluoride sensitivity in S. mutans synergistically reduces C. albicans population in mixed species biofilms

We next examined the effects of a single species’ fluoride sensitivity on the entire 3-species community. For these experiments, we grew surface biofilms for 24 h in the presence of 0.2 mM NaF, and then measured CFUs of each species, total biofilm mass, and media pH. We selected this concentration of fluoride because the ΔCLC^F^ *S. mutans* and ΔFEX *C. albicans* knockout strains exhibit fluoride sensitivity, without being completely eliminated from the biofilm. As a negative control, we performed parallel biofilm experiments in the presence of 0.2 mM NaCl.

Unsurprisingly, in biofilms with a single fluoride-sensitive species, that species was significantly reduced in fluoride-containing media relative to chloride. However, we also observed synergistic reduction in CFUs of the second cariogenic species, especially for biofilms grown with ΔCLC^F^ *S. mutans* (**Figure 3A, Supplementary Figure 2, Supplementary Table 2**). In these three-species biofilms, with WT *C. albicans*, WT *S. gordonii*, and ΔCLC^F^ *S. mutans*, the *S. mutans* population was reduced by a factor of 5×10^2^, and the WT *C. albicans* population by a factor of 7×10^3^ compared to the chloride control. Strikingly, this reduction in the *C. albicans* population upon knockout of the *S. mutans* fluoride exporter is almost as great as the reduction in *C. albicans* population due to knockout of its own fluoride exporter, 5×10^4^-fold. This is consistent with the role of *S. mutans* in extracellular aggregation and biofilm stabilization, which promotes growth of symbionts like *C. albicans*(4, 35). For the three-species biofilms composed of ΔFEX *C. albicans,* WT *S. mutans*, and WT *S. gordonii*, in addition to the reduction of ΔFEX *C. albicans* described above, WT *S. mutans* population decreased by a small, but significant factor of 4.4 compared to the chloride control. In both cases, no significant change in the *S. gordonii* population was observed. Only when biofilms were grown with both ΔFEX *C. albicans* and ΔCLC^F^ *S. mutans* was there a significant decrease in the *S. gordonii* population compared to the chloride control.

**Figure 3.**
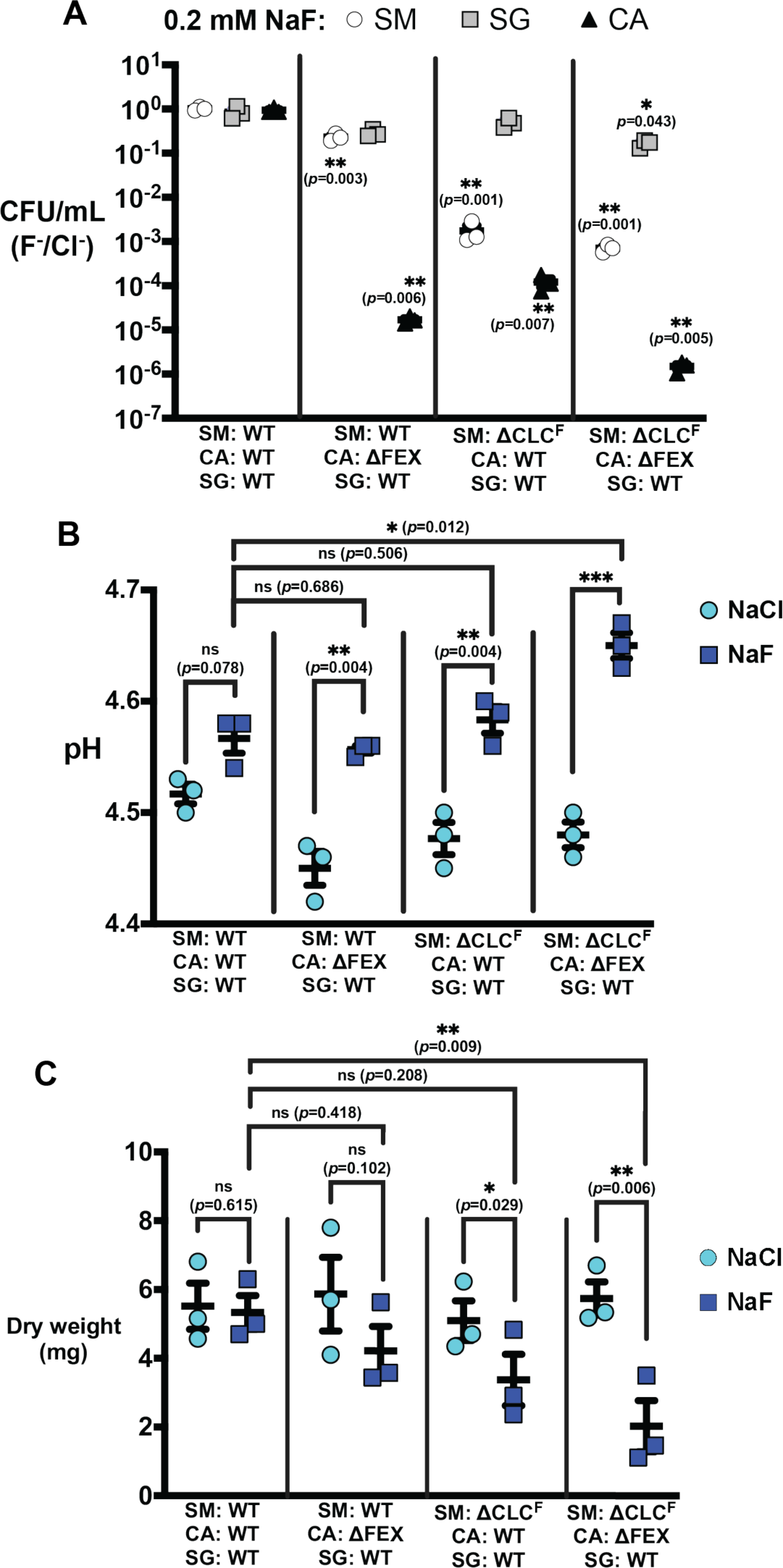
Properties of three-species biofilms grown with ΔCLC^F^ *S. mutans* or ΔFEX *C. albicans* in the presence of 0.2 mM NaF. **A.** CFUs from fluoride-exposed biofilms expressed as a ratio relative to those obtained from control chloride-exposed biofilms. Unnormalized counts from the fluoride-treated and control biofilms are shown in **Supplementary Figure 2**. Significance (p<0.05) shown relative to the ratio of CFUs obtained from the all-WT biofilms. **B.** Media pH after 24 h biofilm growth. **C.** Biofilm mass (dry weight) after 24 h growth. For all panels, datapoints represent mean and SEM of three independent experiments (*n*=3). Significance was calculated using two-way analysis of variance (ANOVA) followed by Fisher’s LSD test. Statistical significance is represented as ‘*’ (*p*< 0.05), ‘**’ (*p*< 0.01) and ‘***’ (*p*< 0.001).

For all biofilms with ΔFEX *C. albicans,* ΔCLC^F^ *S. mutans*, or both, fluoride treatment significantly increased the pH of the biofilm media relative to the NaCl control. However, among the fluoride treated biofilms, only the samples with both ΔCLC^F^ *S. mutans* and ΔFEX *C. albicans* showed significantly higher media pH compared to biofilms with the WT counterparts (**Figure 3B)**. Both biofilms with ΔCLC^F^ *S. mutans* exhibited significantly lower dry weight biomass compared to all-WT or chloride-treated controls, in accord with *S. mutans*’ role as a keystone pathogen whose fitness impacts biofilm stability and assembly (**Figure 3C**). Together these experiments demonstrate that fluoride sensitivity in pathogens, especially *S. mutans*, impact biofilm species composition, mass, and acidogenicity.

### The importance of fluoride export to fitness is recapitulated in biofilms exposed to feast-and-famine cycles

The experiments described above suggest that biofilms composed of fluoride-sensitive strains will be less pathogenic than WT biofilms. To evaluate this in a more faithful model of dental caries, we next grew biofilms on hydroxyapatite, the main chemical component of tooth enamel. For these experiments, we exposed the biofilms to feast-and-famine periods that mimic the cycles of daytime sucrose consumption and overnight fasting experienced by oral microbiota (**Figure 4A)**. We exposed the biofilms to the same sub-lethal concentration of 0.2 mM NaF (or 0.2 mM NaCl as a control) after the biofilm was established, beginning on day 2, and examined the biofilm species composition after each feast and famine period for 5 days (**Figures 4B-D**). For all-WT biofilms, the counts of *S. mutans* and *C. albicans* both increased over the course of the experiment, reaching a steady state by day 5. In contrast, *S. gordonii* counts peaked on day 3, before being outcompeted by the other two species. When biofilms were prepared with ΔFEX *C. albicans*, the overall trend of *S. mutans* and *S. gordonii* growth was preserved, along with the expected reduction in ΔFEX *C. albicans* over the course of the experiment. However, for biofilms prepared with ΔCLC^F^ *S. mutans*, populations of other species were impacted as well. Although the *C. albicans* population increased over the course of the experiment, there was a small reduction in total counts relative to its growth in chloride-exposed biofilms (*p*-values between 0.031 and 0.221 at each timepoint beyond day 2). The *S. gordonii* population also increased over the course of the experiment, reaching a similar population at day 5 as in the all-WT or ΔFEX *C. albicans* experiments, but without the population peak and collapse observed in those two samples. A similar trend in *S. gordonii* growth was also present in the biofilm sample with both ΔFEX *C. albicans* and ΔCLC^F^ *S. mutans*. As in the surface biofilms, synergy between *S. mutans* and *C. albicans* growth was observed, as counts for both species were significantly reduced in biofilms with both knockout strains, compared with biofilms with either individual knockout strain (**Figures 4B-E, Supplementary Fig. 3**).

**Figure 4.**
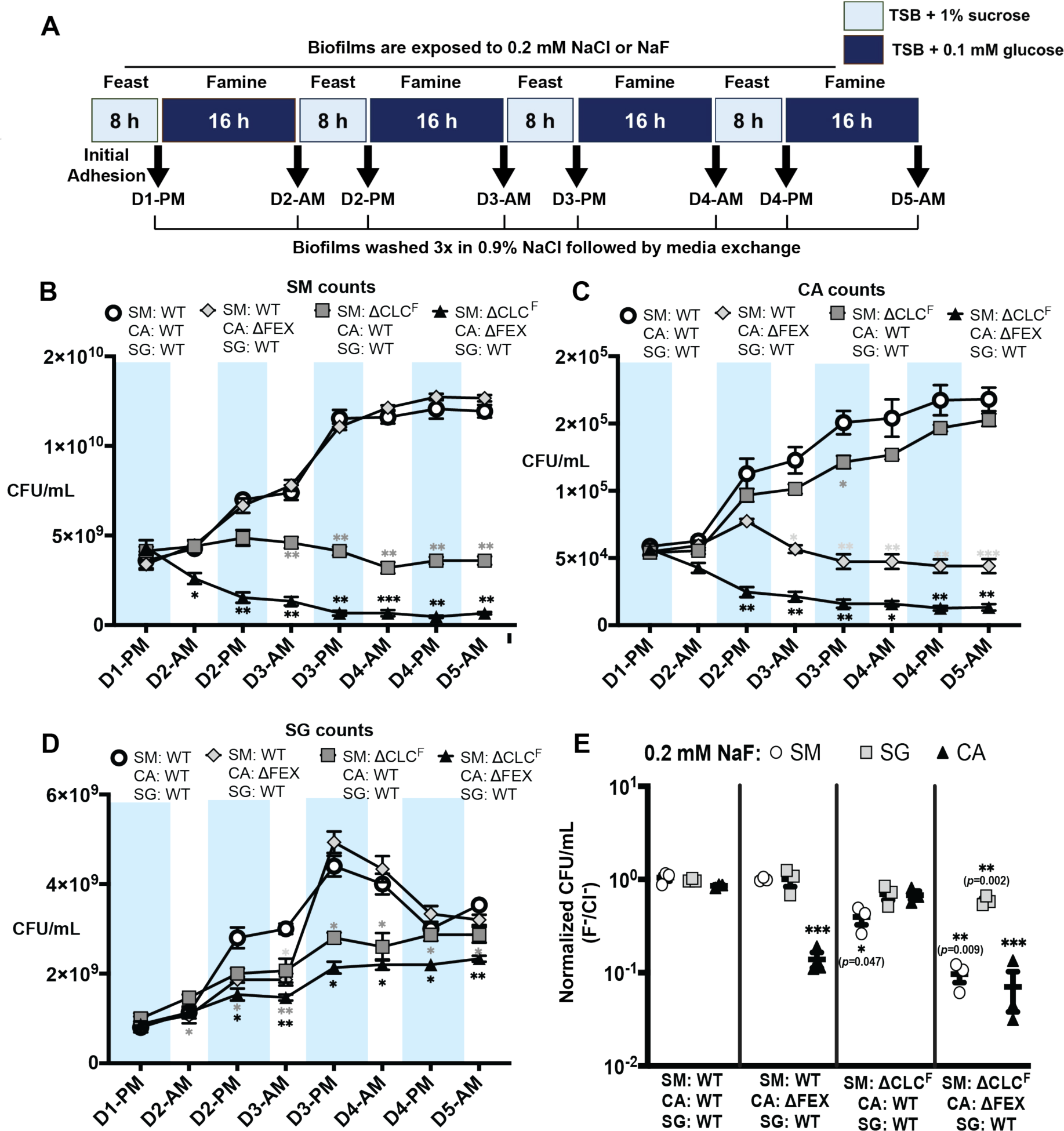
Species composition of biofilms grown on hydroxyapatite discs with feast/famine cycles. **A.** Schematic of media changes for the feast-and-famine dental biofilm model. **B.** Final counts of CFUs for each microbial species at the end of the 5-day experiment. CFUs for biofilms grown in the presence of 0.2 mM NaF are normalized relative to biofilms grown in the presence of 0.2 mM NaCl. Unnormalized counts from the fluoride-treated and control biofilms are shown in **Supplementary Figure 3**. **C.** *S. mutans* CFUs recovered from biofilms of the indicated composition after each treatment in the feast and famine cycle. Statistical significance is shown relative to all-WT fluoride-treated biofilms (open circles). **D.** *C. albicans* CFUs recovered from biofilms of the indicated composition after each treatment in the feast and famine cycle. Statistical significance is shown relative to all-WT biofilms, fluoride-treated biofilms (open circles). **E.** *S. gordonii* CFUs recovered from biofilms of the indicated composition after each treatment in the feast and famine cycle. Statistical significance is shown relative to all-WT biofilms, fluoride-treated biofilms (open circles). For all panels, datapoints represent mean and SEM of three independent experiments (*n*=3). Significance was calculated using two-way analysis of variance (ANOVA) followed by Fisher’s LSD test. Statistical significance is shown relative to parallel experiments in the presence of 0.2 mm NaCl, and is represented as ‘*’ (*p*< 0.05), ‘**’ (*p*< 0.01) and ‘***’ (*p*< 0.001).

### Fluoride sensitivity in S. mutans or C. albicans reduces the pathogenic potential of the biofilm

We also examined several indicators of disease progression over the course of the multiday experiment, including media pH and calcium release from the hydroxyapatite (**Figures 5A, B**), and, at the end of the experiment, biofilm mass and biofilm fluoride content (**Figures 5C, D**). These measurements show pH oscillations characteristic of the feast and famine periods. Drastic reductions in pH occur during every feast period due to sucrose fermentation, and much smaller reductions in pH are seen during the overnight famine period, because the limiting glucose concentration minimizes fermentation. The pH at the end of the feast period was consistently higher throughout the experiment for fluoride-treated biofilms in which either of the pathogenic microorganisms was fluoride export-deficient (**Figure 5A**, **Supplementary Table 3**). The lowest acid production was observed in the fluoride-treated biofilms grown with mutant strains of both *S. mutans* and *C. albicans*. A significant reduction in acid production by the fluoride-treated biofilms prepared with both fluoride-sensitive species persisted even after famine periods, despite the shift away from sugar fermentation.

**Figure 5.**
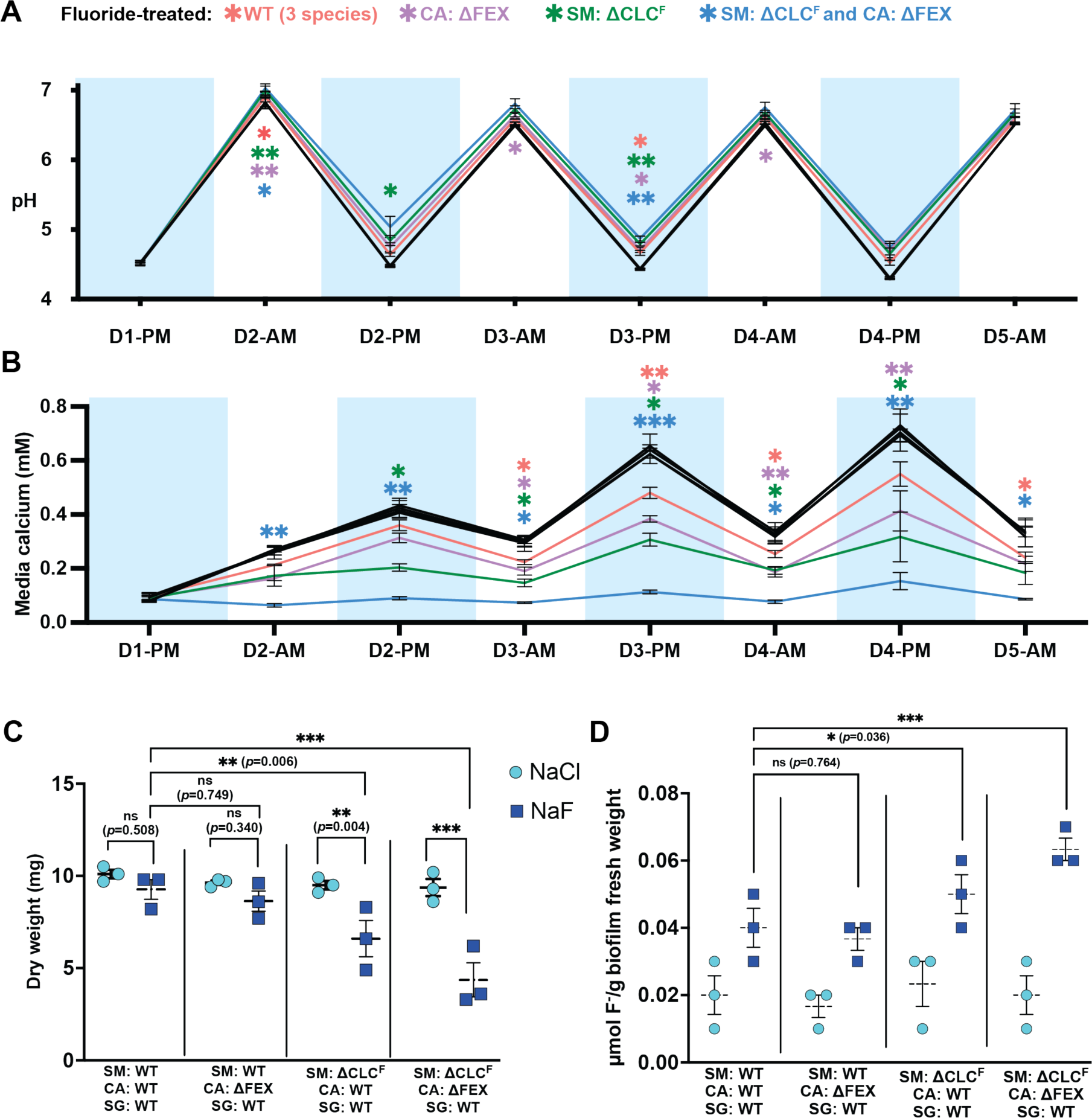
Properties of biofilms grown on hydroxyapatite discs with feast/famine cycles. **A.** Media pH measured after each cycle of feast and famine. **B.** Calcium concentration in media measured after each cycle of feast and famine. For panels A and B, for each biofilm composition, control experiments were performed in the presence of NaCl (black points/lines in panels A and B). These show close correspondence with overlapping points. For the fluoride-treated conditions, statistical significance is shown relative to the chloride-treated biofilms, and is represented as ‘*’ (*p*< 0.05), ‘**’ (*p*< 0.01) and ‘***’ (*p*< 0.001).**C.** Final biofilm mass at the end of the 5-day experiment. **D.** Fluoride content of biofilm at the end of the 5-day experiment. For all panels, datapoints represent mean and SEM of three independent experiments (*n*=3). Significance was calculated using two-way analysis of variance (ANOVA) followed by Fisher’s LSD test.

Similar to pH, we observed oscillations in the media calcium content corresponding to the periods of feast and famine, since acid produced during sucrose fermentation accelerates dissolution of the hydroxyapatite discs (36). Fluoride helps prevent demineralization, so even all-WT biofilms showed lower calcium release into the media compared to the chloride-treated biofilms. However, the biofilms prepared with fluoride-sensitive strains exhibited even lower levels of hydroxyapatite dissolution. Temporal release of calcium was the lowest in the fluoride-treated biofilms grown from combination of the mutant strains of both pathogens, with statistically significant differences compared to the chloride-treated biofilms emerging at all timepoints after fluoride addition, which occurred after the famine treatment on the second day (D2-AM) (**Figure 5B**). The biofilms with ΔCLC^F^ *S. mutans* or ΔFEX *C. albicans* also exhibited significantly lower hydroxyapatite dissolution at most timepoints.

As in the surface biofilms, the biomass of the mature biofilm was significantly reduced for fluoride-treated biofilms prepared with ΔCLC^F^ *S. mutans*, compared to the respective chloride-treated biofilms (**Figure 5C)**. Interestingly, these biofilms also accumulate the highest levels of fluoride (**Figure 5D**). We propose that this is due to fluoride accumulation in the cytoplasm of the species that lack a fluoride exporter, as has been shown to occur in other fluoride export-deficient bacteria (17, 20). When the fluoride concentration is increased to the level that causes lysis and elimination of *S. mutans*, 0.3 mM, fluoride accumulation in the biofilm is likewise eliminated (**Supplementary Figure 4**).

### Only one open reading frame contributes to the functional CLC^F^ exporter in S. mutans

The experiments thus far suggest that the *S. mutans* CLC^F^ is essential to the stability of the cariogenic dental biofilm under conditions of fluoride exposure. Thus, we sought to determine whether ORF 1289, ORF 1290, or both genes encode the functional, fluoride-exporting unit. Across kingdoms of life, CLC proteins form functional homodimers (37), but we also considered the possibility that the *S. mutans* CLC^F^ might be a heterodimer of both gene products. While the contribution of both genes to fluoride resistance has been examined for *S. mutans* UA159 previously, different studies have come to different conclusions about whether 1289 only (19), or both 1289 and 1290 (26), contribute to fluoride efflux.

We first performed bacterial dilution assays for *S. mutans* strains with ORF 1289 and ORF 1290 knocked out individually (referred to as Δ1289 and Δ1290, respectively). Δ1289 *S. mutans* exhibited comparable fluoride sensitivity to the ΔCLC^F^ *S. mutans* strain used for earlier experiments, in which both open reading frames are deleted. In contrast, growth of Δ1290 *S. mutans* was not affected at 0.1 mM NaF, similar to the WT strain, and showed only a marginal sensitivity to 0.3 mM NaF (**Figure 6A**). These data suggest that ORF 1289 contributes more towards the fluoride resistance phenotype.

**Figure 6.**
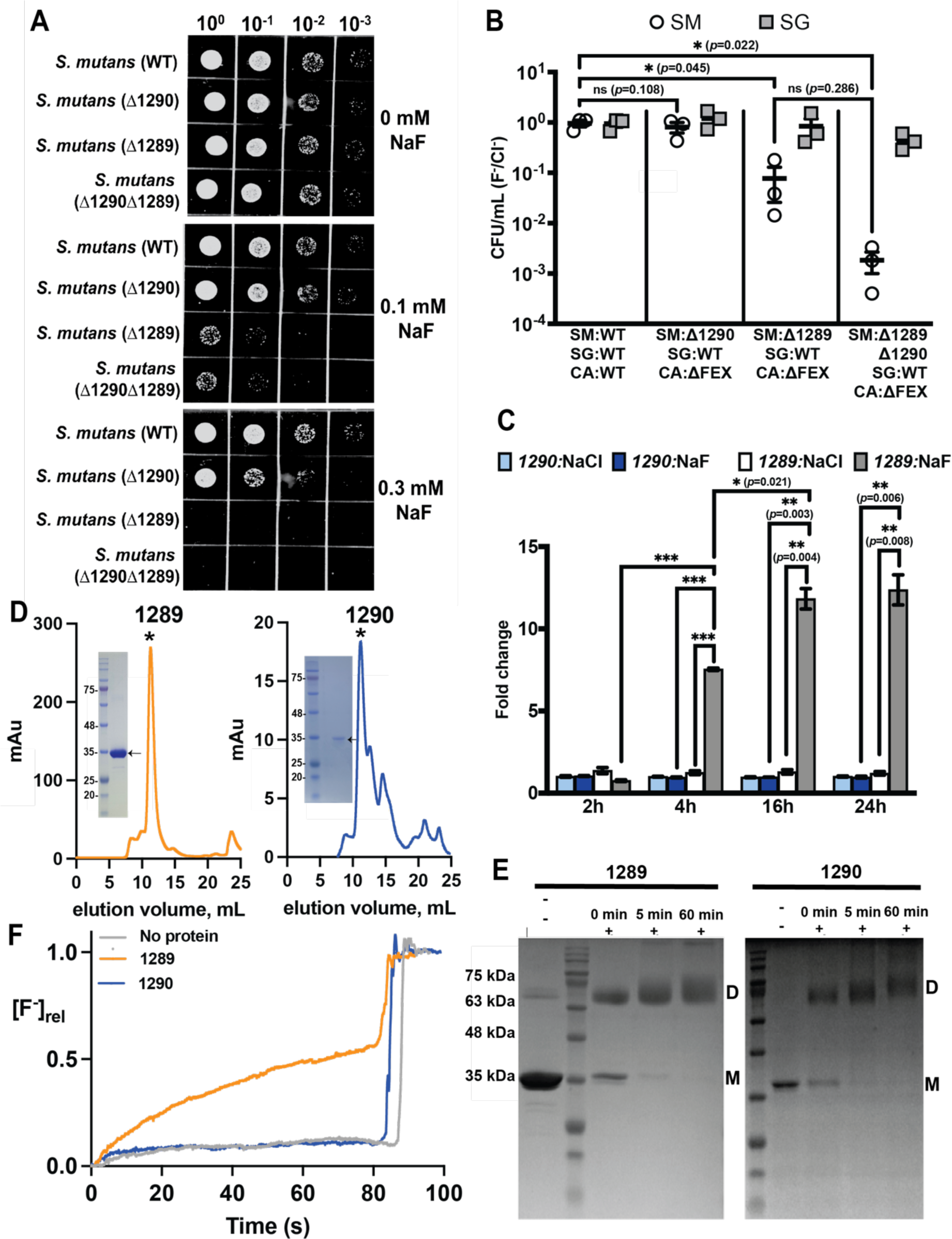
Contributions of ORF 1289 and ORF 1290 to fluoride resistance and transport in *S. mutans*. **A.** 10-fold serial dilutions of Δ1289 *S. mutans*, Δ1290 *S. mutans,* and the ΔCLC^F^ strain (Δ1290Δ1289) used in prior experiments. Plates contain NaF as indicated. **B.** CFUs of *S. mutans* and *S. gordonii* harvested from three-species (*S. mutans, S. gordonii, C. albicans*) biofilms after 24 h growth in the presence of 0.2 mM NaF, normalized relative to CFUs harvested from biofilms grown in the presence of 0.2 mM NaCl. Significance is expressed relative to the corresponding biofilm grown with 0.2 mM NaCl, and was calculated using two-way analysis of variance (ANOVA) followed by Fisher’s LSD test. **C.** Gene expression profile of *S. mutans* ORFs 1290 and 1289 in 3-species biofilms (WT *S. mutans*) grown for variable lengths of time. Gene expression is shown as fold-change relative to housekeeping gene *gyrase A*. Datapoints represent mean and SEM of three independent biological replicates (*n*=3). Significance was calculated using two-way analysis of variance (ANOVA) followed by Fisher’s LSD test. **D.** Size exclusion chromatograms (SEC) of purified proteins encoded by ORF 1289 (left) and ORF 1290 (right). The asterisks indicate the peak that was collected and reconstituted for functional analysis. Insets: SDS-PAGE gel of major SEC peak stained with Coomassie. Note that membrane proteins often run at a lower molecular weight than expected due to incomplete denaturation in SDS detergent. **E.** Crosslinking analysis of proteins from panel D. Samples were exposed to 0.125% glutaraldehyde for the indicated time. Bands consistent with a monomer and dimer are indicated by the labels “M” and “D,” respectively. **F.** Fluoride efflux from liposomes reconstituted with 1289 (orange), 1290 (blue), or no protein (gray). Fluoride transport is initiated by valinomycin addition at 5 s. Traces are normalized relative to total encapsulated F^−^, which is determined after fluoride addition (sharp increase in F^−^ at ∼80 s.). Traces are representative of data from three independent proteoliposome preparations. The rate of transport was 830±20 F^−^/s for ORF 1289 and 17±5 F^−^/s for ORF 1290 (mean and SEM from three liposome preparations).

We next tested the hypothesis that ORF 1290 is expressed at a different timepoint or under different nutrient conditions inherent to biofilm growth. We examined Δ1289, Δ1290, and the double knockout in three species surface biofilms with *S. gordonii* and *C. albicans.* In agreement with the dilution assays, Δ1289 exhibited significantly reduced survival in fluoride compared to chloride, comparable to the ΔCLC^F^ strain with deletions of both ORFs 1290 and 1289. For biofilms grown with Δ1290, there was a slight, but not statistically significant, decrease in *S. mutans* CFUs harvested from the biofilm (*p* = 0.108; **Figure 6B, Supplementary Fig. 5**). Similarly, the Δ1289 Δ1290 double knockout was not significantly less fit relative to the Δ1289 single knockout in fluoride (p = 0.29; **Figure 6B**).

Quantitative gene expression profiling of samples collected from biofilms formed with WT *S. mutans* corroborated these phenotypic observations. ORF 1289 showed an increase in transcripts over time when the biofilm was treated with 0.2 mM NaF, with a 17-fold increase between 1 and 24 h of fluoride exposure. In contrast, expression of ORF 1290 did not change over 24 h of fluoride treatment, and remained at a similar level as ORFs 1290 and 1289 under chloride treatment (**Figure 6C**).

To assess transport function, we purified the proteins encoded by the individual ORFs and tested their ability to export fluoride upon functional reconstitution in liposomes. Proteins encoded by both ORFs 1289 and 1290 could be heterologously expressed in *E. coli* under a tetracycline promoter and purified. Both proteins showed monodisperse peaks by size exclusion chromatography, an indication of protein stability, although the yield of 1290 was lower (**Figure 6D**). Crosslinking experiments showed that both 1289 and 1290 purify as homodimers, similar to other CLC proteins, providing another indication that both proteins are properly folded and able to assemble (**Figure 6E**). We reconstituted purified 1289 and 1290 in liposomes and monitored fluoride transport by each protein individually using a fluoride electrode-based transport assay(38). 1289 catalyzed robust fluoride efflux from liposomes, with a unitary transport rate of 830±20 ions/second, in line with rates measured for other CLC^F^ transporters (38). In contrast, fluoride efflux from proteoliposomes containing 1290 was indistinguishable from uncatalyzed fluoride leak exhibited by protein-free liposomes (**Figure 6F**). From these lines of evidence – resistance assays, quantitative transcript profiling, and functional reconstitution, we conclude that the functional *S. mutans* CLC^F^ is a homodimer encoded by ORF 1289.

## Discussion

In this work, we sought to understand how fluoride export impacts the fitness of pathogens in dental biofilms, a question with particular relevance to the oral microbiota due to the widespread use of fluoride in oral healthcare products. Moreover, the observation that dental pathogens and oral eubacteria export fluoride using proteins from entirely different molecular families provides an opportunity to specifically target fluoride export by oral pathogens. We therefore examined the effects of fluoride export at the individual species level, in model 3-species surface biofilms, and in biofilms grown on the dental mineral hydroxyapatite under conditions that mimic physiologically realistic cycles of feast and famine.

As expected from other studies of fluoride exporter deletion strains, both *S. mutans* and *C. albicans* are rendered sensitive to hundreds of micromolar F^−^ upon deletion of their respective fluoride exporters. Although biofilms are protective against many antimicrobial stressors, fluoride is inhibitory to a similar extent in a biofilm context. Particularly striking was the elimination of ΔCLC^F^ *S. mutans* from biofilms at a low fluoride concentration of 0.3 mM (⁓5.7 ppm F^−^), a concentration almost 200-fold lower than the fluoride levels in toothpastes and just 8-fold higher than that in fluoridated municipal water supplies (39, 40). Whereas fluoride acts as a bacteriostatic agent for many microbes (18, 20, 41), ΔCLC^F^ *S. mutans* unexpectedly underwent massive lysis under fluoride stress. *S. mutans* is known to induce autolysins in response to stressors such as heat or protein synthesis inhibitors (42). Indeed, partial autolysis of WT *S. mutans* exposed to 5 mM NaF was reported decades ago (43). Typically autolysis is adaptive, eliminating a subpopulation of cells to provide nutrients or genetic material to sister cells. The mechanism of fluoride-induced lysis, and why lysis is not limited to a sub-population, remains to be explored.

Knockout of the *S. mutans* fluoride exporter influences properties of the entire biofilm under fluoride stress. This finding is not unexpected, since *S. mutans* actively contributes to dysbiosis by secreting enzymes that can polymerize sucrose into extracellular polysaccharides, enhancing biofilm mass and providing a substrate for other species growth(44); and further by producing acid that eliminates species that are not tolerant to low pH(45). Biofilms grown with ΔCLC^F^ *S. mutans* exhibited a lower *C. albicans* population, a greater proportion of *S. gordonii*, higher pH, and less demineralization. The total biofilm biomass was also decreased, as fluoride sensitivity in *S. mutans* did not cause overgrowth of *S. gordonii*, which infrequently causes infective endocarditis or other invasive infections (46). While biofilms prepared with ΔFEX *C. albicans* also exhibited lower pathogenicity, the effect was not as large as that of ΔCLC^F^ *S. mutans*.

An unexpected finding was the retention of fluoride in biofilms with ΔCLC^F^ *S. mutans.* This effect mirrors the intracellular fluoride accumulation that has been measured in other microbes that lack fluoride exporters (17, 20, 47). In these cells, intracellular fluoride accumulation quantitatively follows the pH gradient(20). For example, for cells in a pH 5 environment, with a cytoplasmic pH maintained at 7, fluoride will accumulate intracellularly to a concentration 100-fold greater than that of the external fluoride. For ΔCLC^F^ *S. mutans*, we expect that fluoride accumulation only occurs at sub-lethal concentrations; at higher concentrations, we observe cell lysis, which would release fluoride from the biofilm where it can be washed away. This implies that at moderate fluoride concentrations, the bacteria themselves could act as fluoride reservoirs, slowly releasing fluoride into the biofilm over several hours as the population lyses.

Because of its primary importance to *S. mutans* fitness in fluoride-treated biofilms, we sought to determine the functional unit of the *S. mutans* CLC^F^. The literature features contrasting results, with one study concluding that both ORFs 1289 and 1290 contribute to fluoride resistance(26), and another study concluding that 1289 only was responsible for the fluoride resistance phenotype(19) (all studies investigated *S. mutans* UA159). Although the reason for the discrepancy in the previous studies is not clear, we found that 1290 contributed little to resistance, either in culture or in the biofilm context. Moreover, we did not observe fluoride-induced changes in the transcription of ORF 1290, and although we were able to heterologously express and purify a well-folded protein encoded by 1290 under an artificial promoter, it did not transport fluoride in a reconstituted system. 1289, in contrast, is induced by fluoride, and encodes a homodimeric transporter that catalyzes efficient fluoride export from lipid vesicles. Based on the sum of this evidence, we conclude that in *S. mutans* UA159, ORF 1289 is necessary and sufficient for fluoride resistance. Although ORF 1290 presumably originated from a duplication of a functional CLC^F^ gene, it appears that the redundant gene has degraded into a pseudogene that is not expressed or functional. However, we note that it is not obvious from sequence alignments why 1290 is unable to transport fluoride, as all fluoride-binding residues are conserved (**Supplementary Figure 6**). Additionally in a fluoride-resistant derivative of *S. mutans* UA159, mutations in the promoter region 5’ to 1290 contribute to constitutive expression of 1290 and other genes in the operon(48). Thus, we cannot definitively rule out the possibility that in other closely related bacteria from the mutans subgroup, both genes are expressed and contribute to fluoride resistance.

In summary, we report substantial changes in dental biofilm communities based on a single factor, fluoride export (**Figure 7)**. Our results imply that fluoride efflux inhibitors that specifically target the CLC^F^ of *S. mutans* or the FEX channel of *C. albicans* could serve to reduce or eliminate cariogenic species while preserving commensal microbes in the dental biofilm, which mainly export fluoride via the action of the molecularly distinct Fluc channels. Humans also lack known fluoride exporters, including CLC^F^ or FEX proteins, further recommending fluoride efflux as a potential antimicrobial target. The *S. mutans* CLC^F^ is a particularly attractive target for development of such an inhibitor because of its sensitivity to lysis under fluoride stress, its keystone role in biofilm assembly and pathogenesis, and its straightforward architecture as a homodimer encoded by a single open reading frame.

**Fig. 7.**
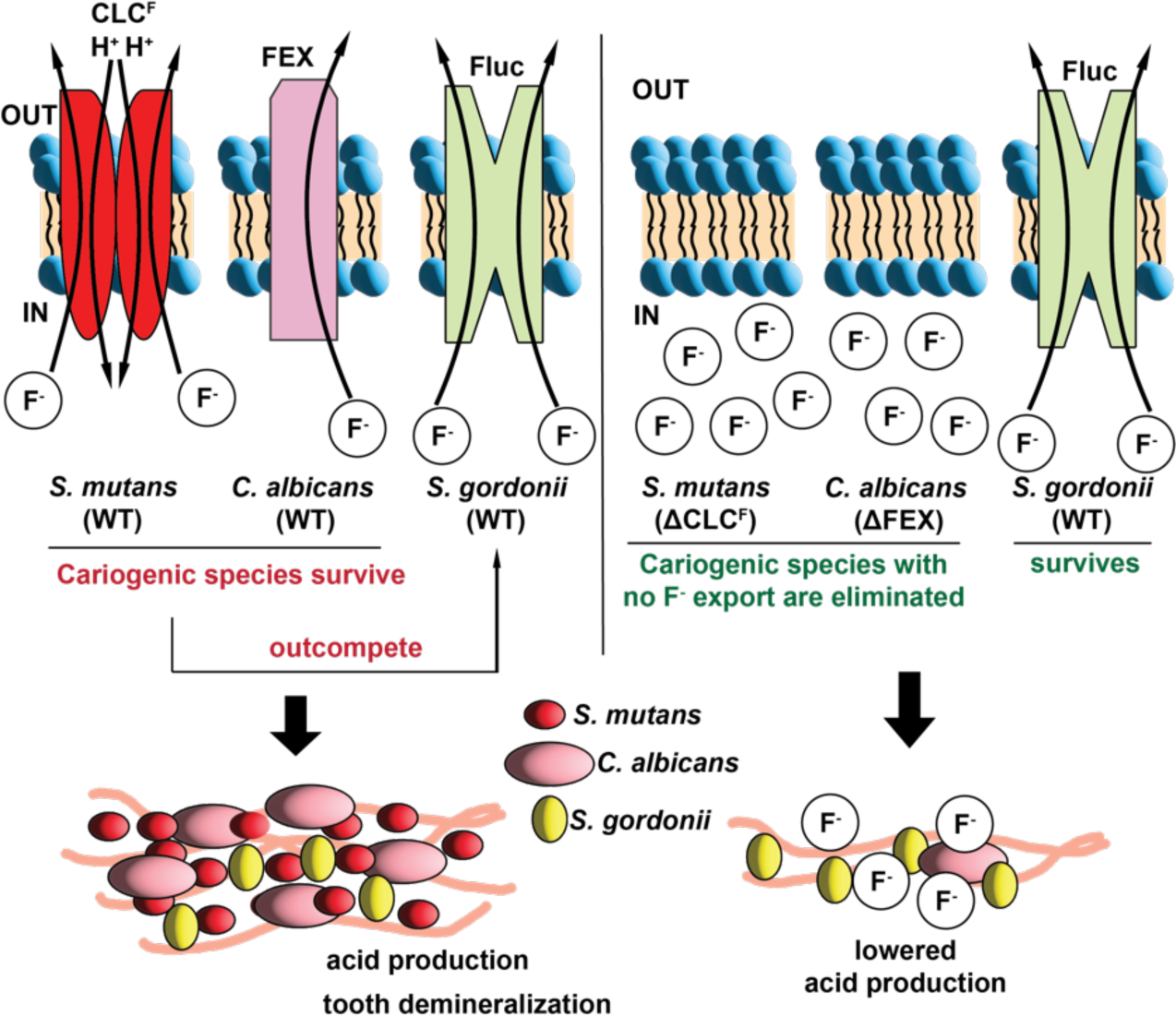
Schematic of species composition in dental biofilms. Left: all WT species. Right: species with deletion of fluoride export gene.

## Acknowledgements

This work was supported by National Institutes of Health grant R21-DE032837 to R.B.S. and L.M.A.T, R21-AI178229 to A.K., and R21-AI168571-01 to A.K. C.-Y. K. was supported by an American Heart Association pre-doctoral fellowship.

## Author Contributions

Aditya Banerjee: conceptualization, investigation, writing – original draft, visualization; Chia-Yu Kang: conceptualization, investigation, visualization; Minjun An: investigation; B. Ben Koff: investigation; Sham Sunder: methodology; Anuj Kumar: methodology, funding acquisition; Livia M. A. Tenuta: conceptualization, writing – review and editing, funding acquisition, project administration; Randy B. Stockbridge: conceptualization, data visualization, writing – review and editing, funding acquisition, project administration.

## Supplementary files

**Supplementary Fig. 1.**
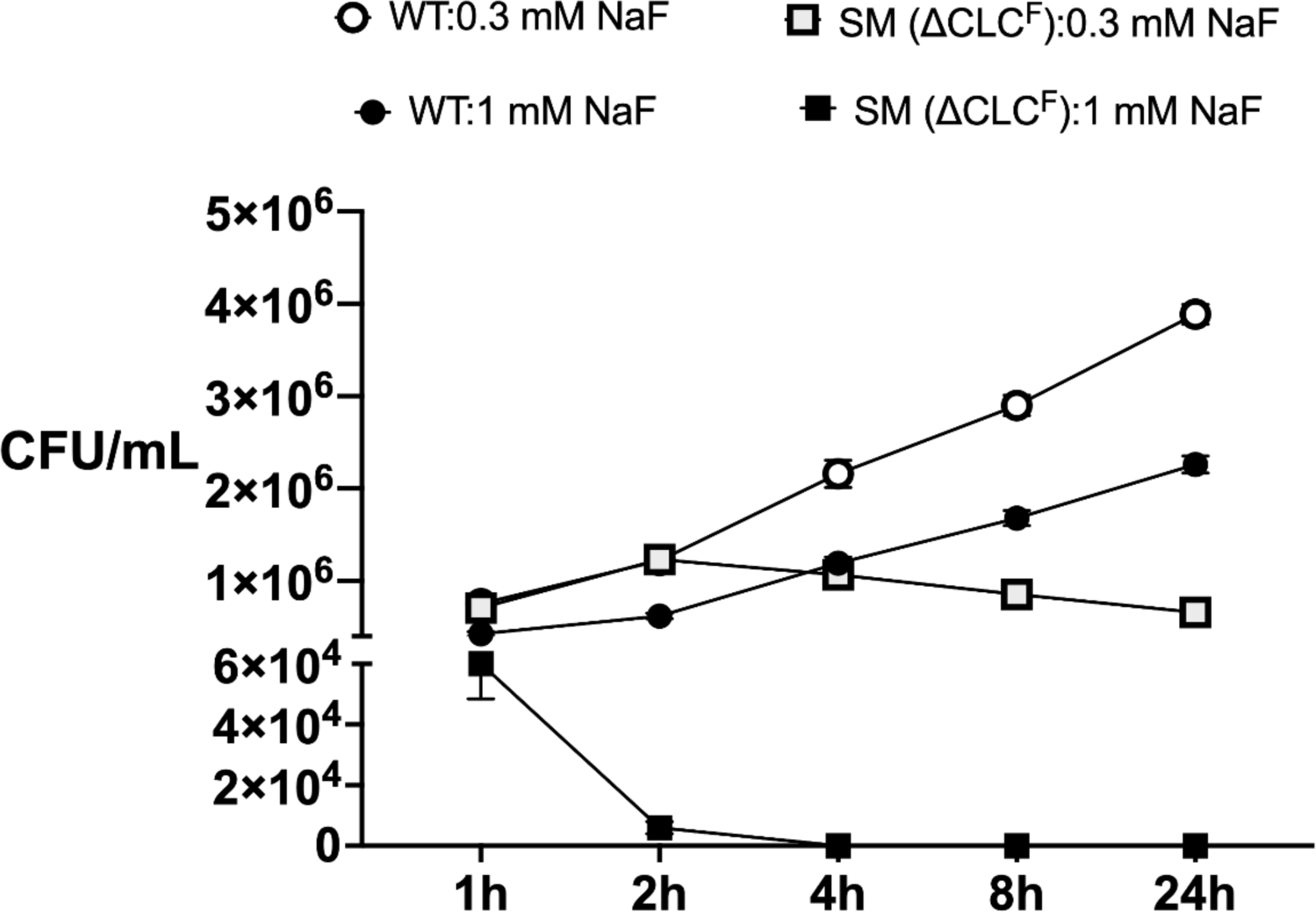
Recovered CFUs of ΔCLC^F^ and WT *S. mutans* as a function of time for samples treated with 0.3 or 1 mM NaF.

**Supplementary Fig. 2.**
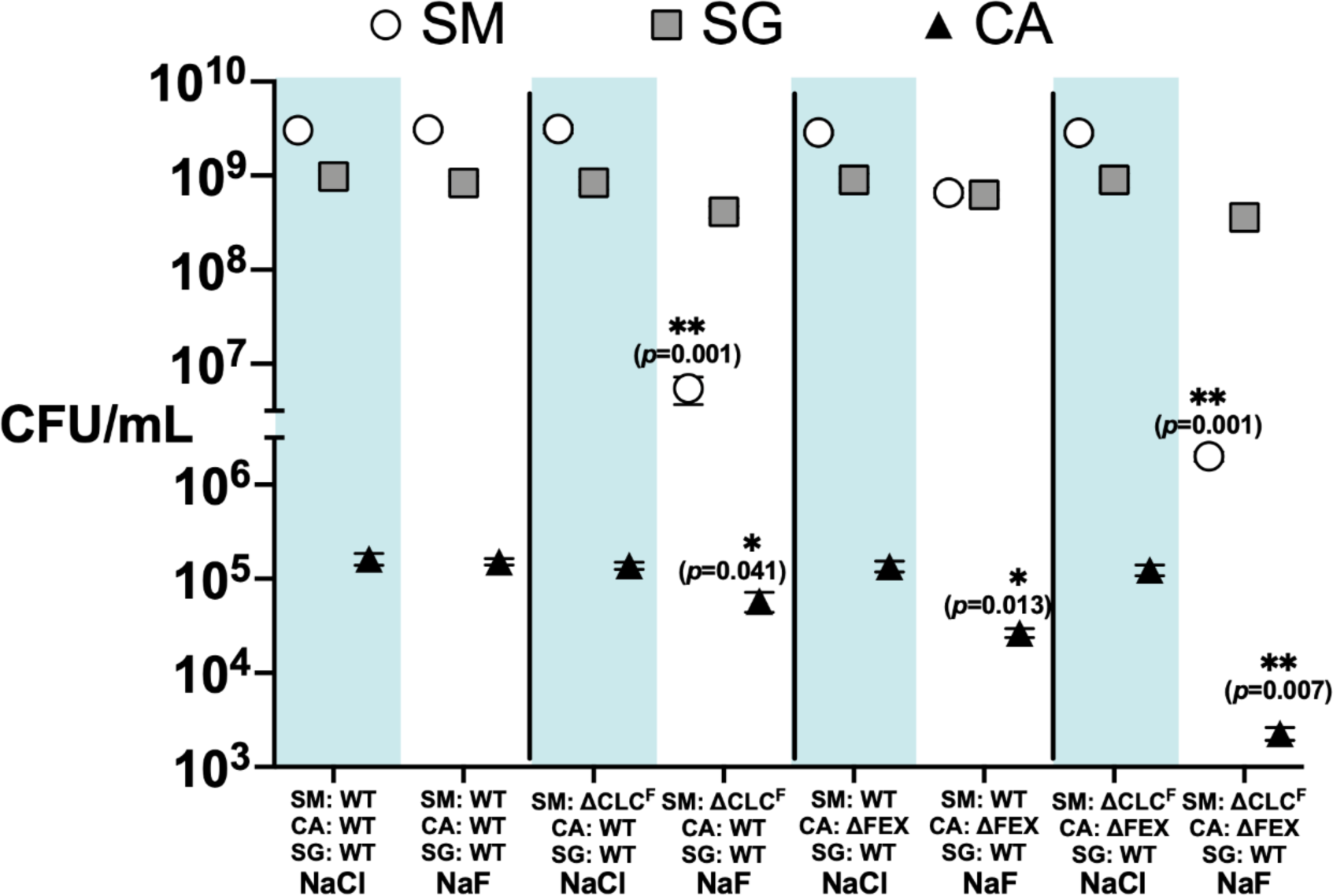
CFUs of *S. mutans*, *S. gordonii* and *C. albicans* recovered from three-species biofilms grown with ΔCLC^F^ *S. mutans* or ΔFEX *C. albicans* or respective WT strains in the presence of 0.2 mM NaCl or NaF, and harvested after 24 h. The datapoints represent mean and SEM of three independent experiments (*n*=3). Significance was calculated using two-way analysis of variance (ANOVA) followed by Fisher’s LSD test. Statistically significant differences (*p*< 0.05) relative to the all-WT biofilms are indicated and represented as ‘*’ (*p*< 0.05), ‘**’ (*p*< 0.01) and ‘***’ (*p*< 0.001).

**Supplementary Fig. 3.**
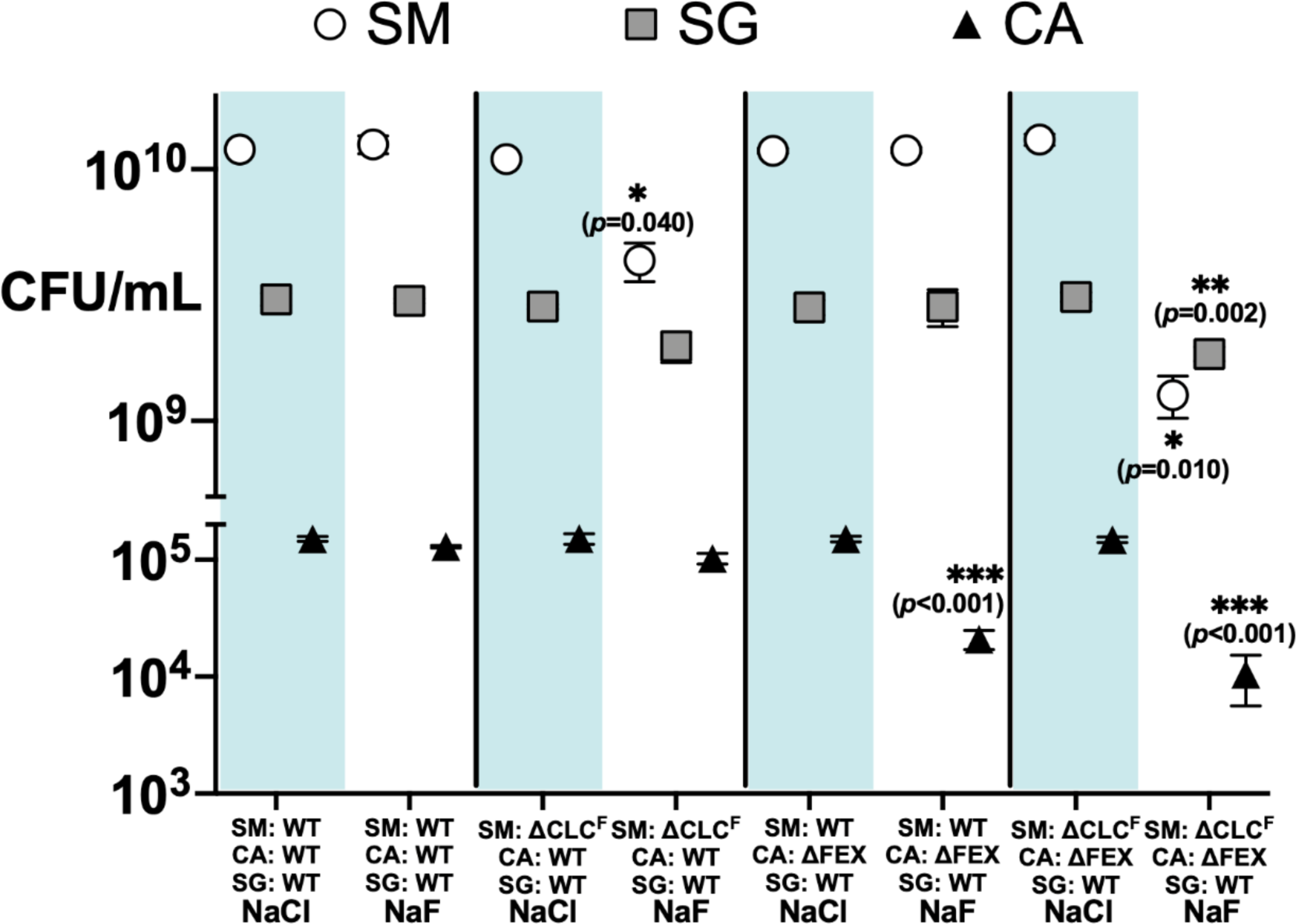
Final counts of CFUs for *S. mutans*, *S. gordonii* and *C. albicans* recovered from three-species biofilms grown with ΔCLC^F^ *S. mutans* or ΔFEX *C. albicans* or respective WT strains in the presence of 0.2 mM NaCl or NaF and at the end of the 5-day feast-and-famine experiment. The datapoints represent mean and SEM of three independent experiments (*n*=3). Significance was calculated using two-way analysis of variance (ANOVA) followed by Fisher’s LSD test. Statistical significance is represented as ‘*’ (*p*< 0.05), ‘**’ (*p*< 0.01) and ‘***’ (*p*< 0.001).

**Supplementary Fig. 4.**
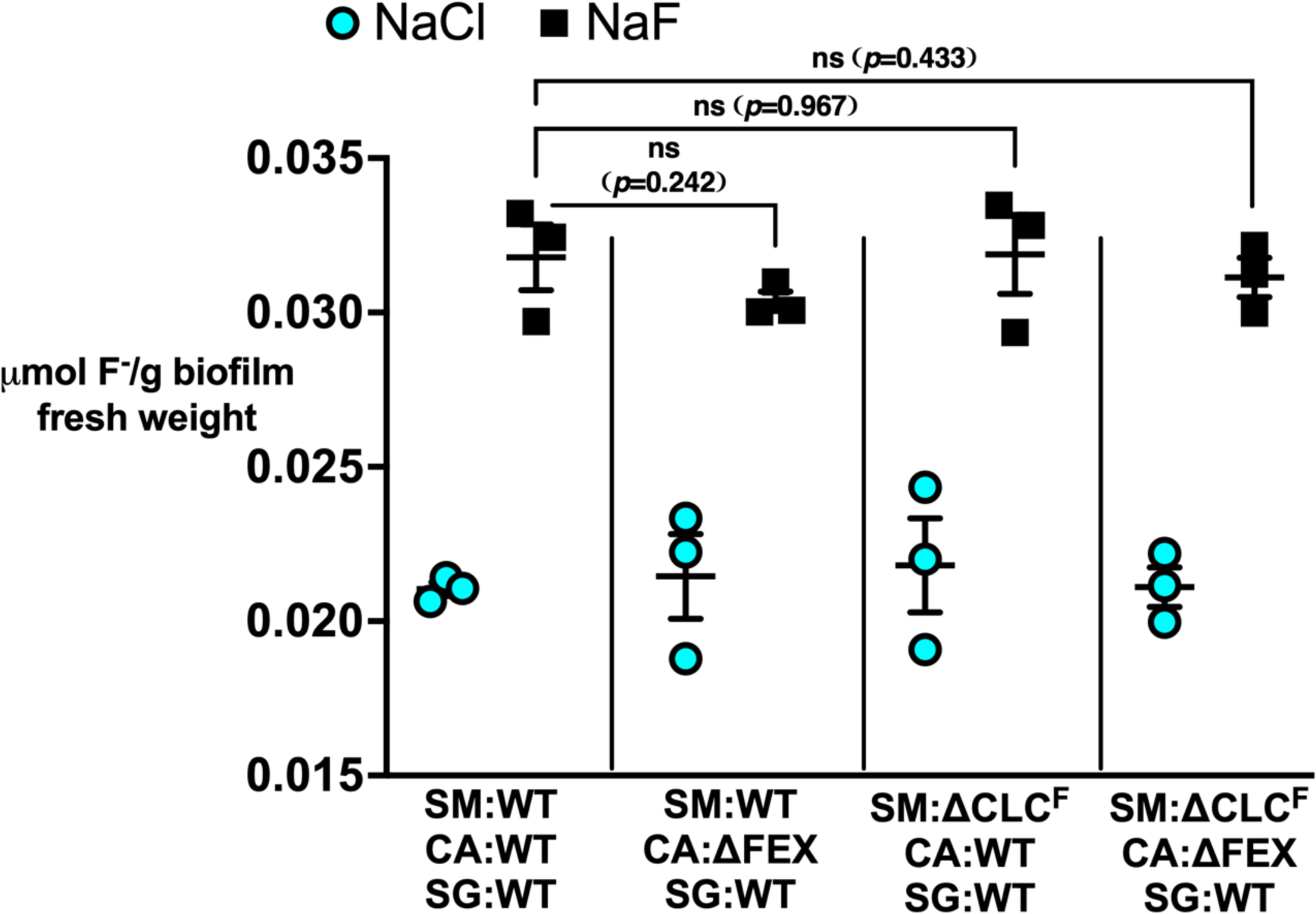
Fluoride content of biofilm combinations exposed to 0.3 mM NaCl or NaF and at the end of the 5-day experiment. For all panels, datapoints represent mean and SEM of three independent experiments (*n*=3). Significance was calculated using two-way analysis of variance (ANOVA) followed by Fisher’s LSD test and is represented as ‘*’ (*p*< 0.05).

**Supplementary Fig. 5.**
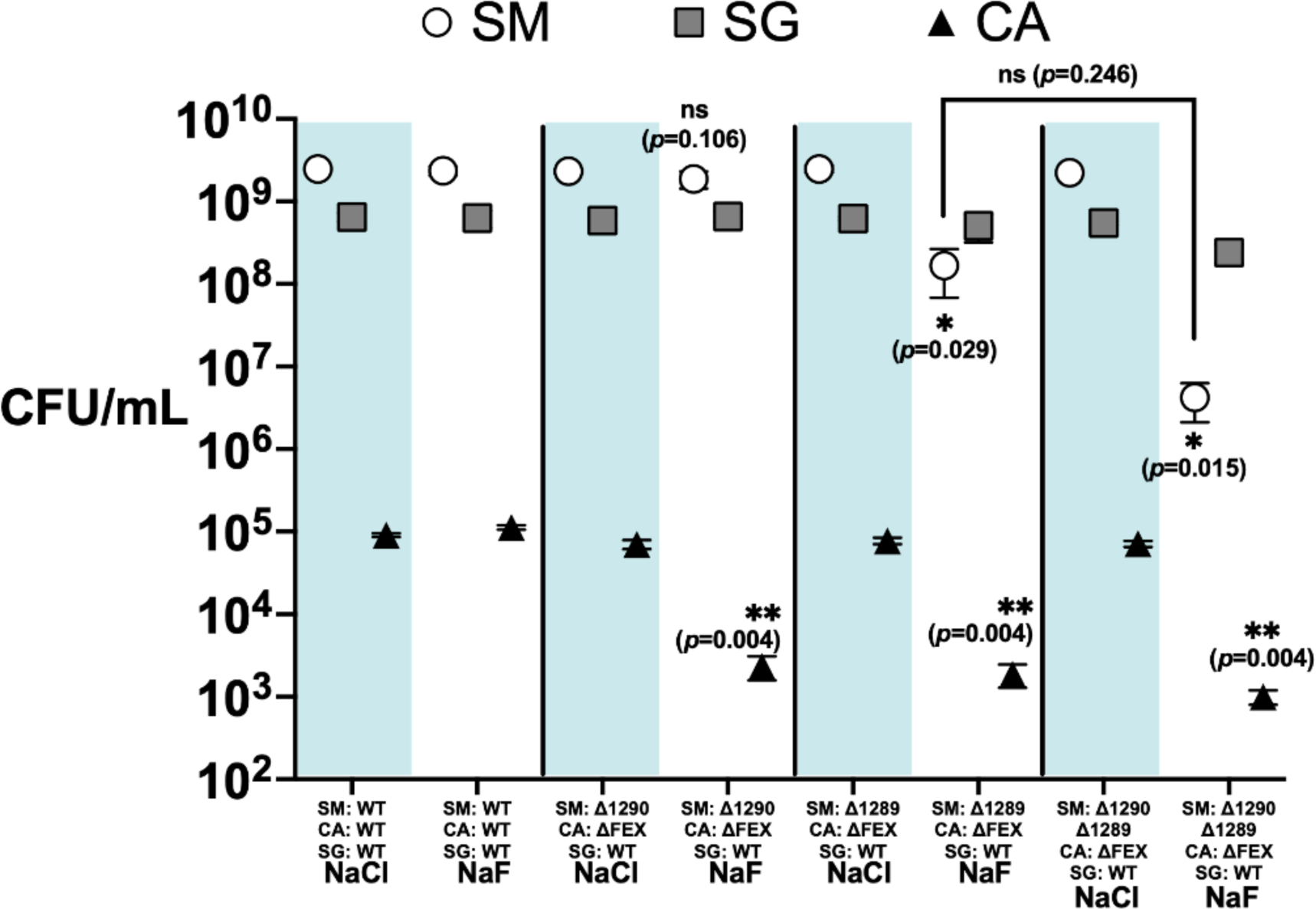
CFUs of *S. mutans*, *S. gordonii*, and *C. albicans* harvested from three-species (*S. mutans, S. gordonii, C. albicans*) biofilms derived from Δ1290 or Δ1289 or Δ1290Δ1289 *S. mutans*, ΔFEX *C. albicans* or the respective WT strains after 24 h growth in the presence of 0.2 mM NaCl or NaF. The datapoints represent mean and SEM of three independent experiments (*n*=3). Statistical significance relative to the all-WT biofilms is shown. Significance was calculated using two-way analysis of variance (ANOVA) followed by Fisher’s LSD test. Statistical significance is represented as ‘*’ (*p*< 0.05), ‘**’ (*p*< 0.01) and ‘***’ (*p*< 0.001).

**Supplementary Fig. 6.**
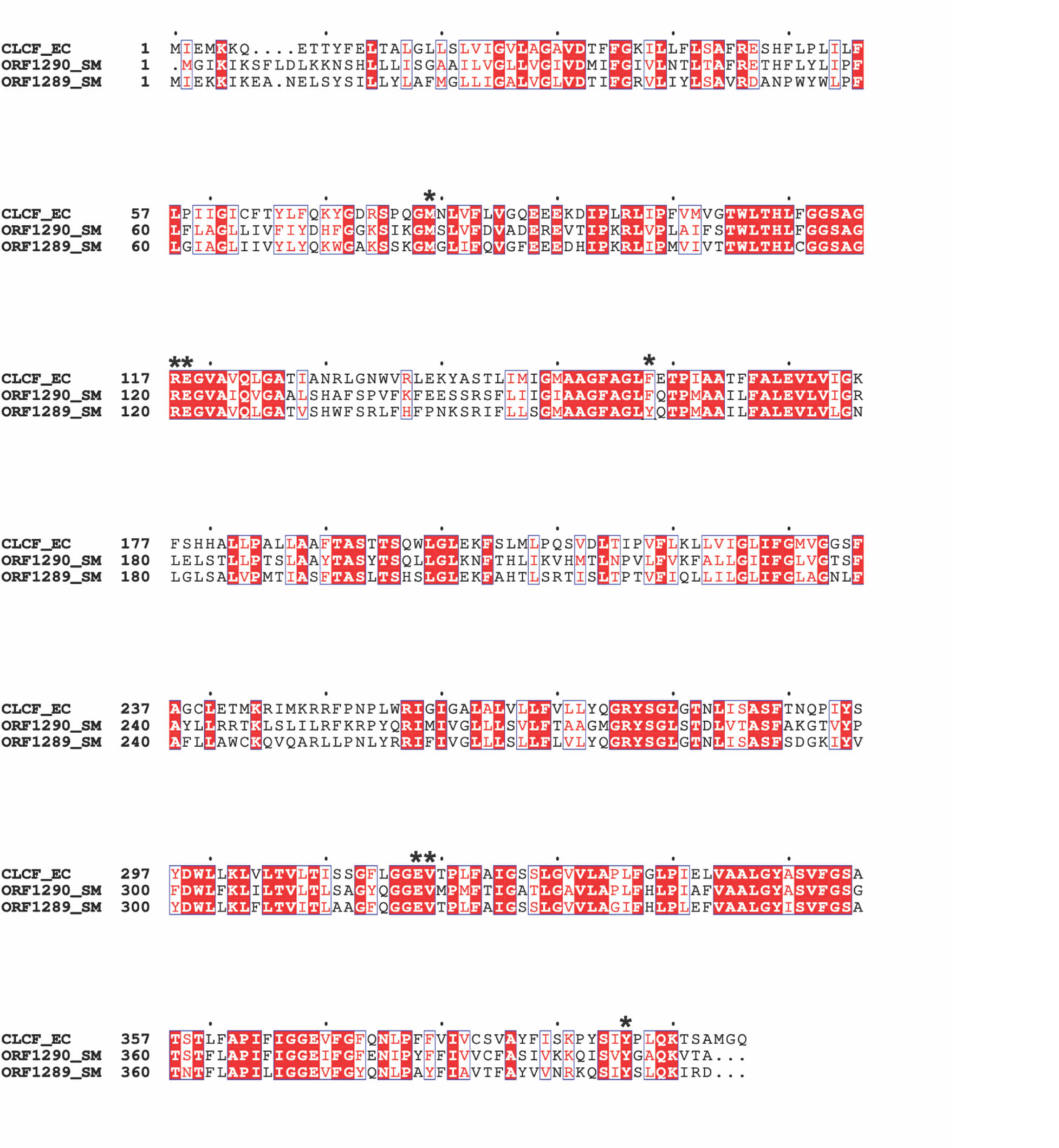
Multiple sequence alignment of *Enterococcus casseliflavus* CLC^F^ (GenBank EEV30821.1), *S. mutans* ORF 1290 (NCBI Protein ID: AAN58967.1) and *S. mutans* ORF 1289 (NCBI Protein ID: AAN58966. Residues that have been functionally implicated in fluoride binding and transport are marked with asterisks.

**Supplementary Table 1:**
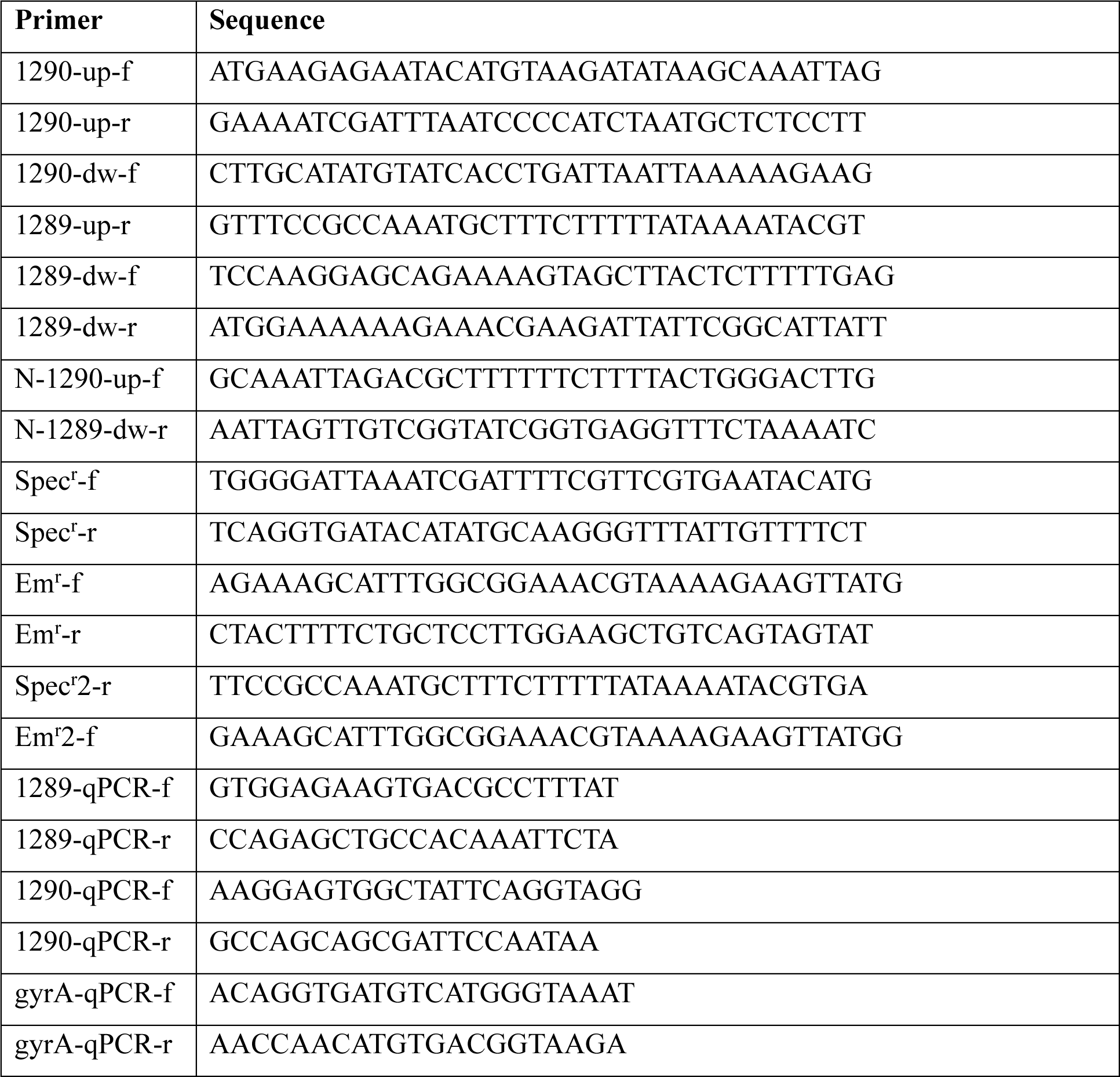
List of primers used in the study.

**Supplementary Table 2:** Statistical significance table for microbial quantification in polystyrene stalks. Significance was calculated using two-way analysis of variance (ANOVA) followed by Fisher’s LSD test implemented in GraphPad Prism 8. Statistical significance is represented as ‘*’ (*p*< 0.05), ‘**’ (*p*< 0.01) and ‘***’ (*p*< 0.001).

| Organism | Combination | Treatments | Summary | Adjusted 'p' Value |
| --- | --- | --- | --- | --- |
| Microbial quantification of biofilm formed with single or both mutant strains of *S. mutans* or *C. albicans* |  |  |  |  |
| *S. mutans* | WT (3 species) | NaCl vs. NaF | ns | 0.8126 |
| | SM (ΔCLC<sup>F</sup>) | NaCl vs. NaF | * | 0.0103 |
| | CA (ΔFEX) | NaCl vs. NaF | ** | 0.0021 |
| | SM (ΔCLC<sup>F</sup>)-CA (ΔFEX) | NaCl vs. NaF | ** | 0.004 |
| | WT (3 species) vs. SM (ΔCLC<sup>F</sup>) | NaF | ** | 0.0016 |
| | WT (3 species) vs. CA (ΔFEX) | NaF | ** | 0.0032 |
| | WT (3 species) vs. SM (ΔCLC<sup>F</sup>)-CA (ΔFEX) | NaF | ** | 0.0015 |
| *C. albicans* | WT (3 species) | NaCl vs. NaF | ns | 0.7473 |
| | SM (ΔCLC<sup>F</sup>) | NaCl vs. NaF | ns | 0.0828 |
| | CA (ΔFEX) | NaCl vs. NaF | * | 0.0184 |
| | SM (ΔCLC<sup>F</sup>)-CA (ΔFEX) | NaCl vs. NaF | * | 0.0167 |
| | WT (3 species) vs. SM (ΔCLC<sup>F</sup>) | NaF | * | 0.0412 |
| | WT (3 species) vs. CA (ΔFEX) | NaF | * | 0.013 |
| | WT (3 species) vs. SM (ΔCLC<sup>F</sup>)-CA (ΔFEX) | NaF | ** | 0.0067 |
| *S. gordonii* | WT (3 species) | NaCl vs. NaF | ns | 0.821 |
| | SM (ΔCLC<sup>F</sup>) | NaCl vs. NaF | ns | 0.1212 |
| | CA (ΔFEX) | NaCl vs. NaF | ns | 0.0821 |
| | SM (ΔCLC<sup>F</sup>)-CA (ΔFEX) | NaCl vs. NaF | ns | 0.0671 |
| | WT (3 species) vs. SM (ΔCLC<sup>F</sup>) | NaF | ns | 0.182 |
| | WT (3 species) vs. CA (ΔFEX) | NaF | ns | 0.3416 |
| | WT (3 species) vs. SM (ΔCLC<sup>F</sup>)-CA (ΔFEX) | NaF | ns | 0.0767 |
| Microbial quantification of biofilm formed with strains of *S. mutans* with either or both the CLC<sup>F</sup> ORFs knocked out, *C. albicans* ΔFEX and *S. gordonii* WT | | | | |
| *S. mutans* | WT (3 species) | NaCl vs. NaF | ns | 0.7687 |
| | SM (Δ1290)- | NaCl vs. NaF | ns | 0.4119 |
| | CA (ΔFEX) | | | |
| | SM (Δ1289)-<br>CA (ΔFEX) | NaCl vs. NaF | * | 0.0211 |
| | SM (Δ1290Δ1289)-<br>CA (ΔFEX) | NaCl vs. NaF | ** | 0.0019 |
| | WT (3 species) vs.<br>SM (Δ1290)-<br>CA (ΔFEX) | NaF | ns | 0.1060 |
| | WT (3 species) vs.<br>SM (Δ1289)-<br>CA (ΔFEX) | NaF | * | 0.0291 |
| | WT (3 species) vs.<br>SM (Δ1290Δ1289)-<br>CA (ΔFEX) | NaF | * | 0.0151 |
| | SM (Δ1289)-<br>CA (ΔFEX) vs.<br>SM (Δ1290Δ1289)-<br>CA (ΔFEX) | NaF | ns | 0.2460 |
| *C. albicans* | WT (3 species) | NaCl vs. NaF | ns | 0.0533 |
| | SM (Δ1290)-<br>CA (ΔFEX) | NaCl vs. NaF | * | 0.0178 |
| | SM (Δ1289)-<br>CA (ΔFEX) | NaCl vs. NaF | * | 0.0100 |
| | SM (Δ1290Δ1289)-<br>CA (ΔFEX) | NaCl vs. NaF | ** | 0.0066 |
| | WT (3 species) vs.<br>SM (Δ1290)-<br>CA (ΔFEX) | NaF | ** | 0.0037 |
| | WT (3 species) vs.<br>SM (Δ1289)-<br>CA (ΔFEX) | NaF | ** | 0.0037 |
| | WT (3 species) vs.<br>SM (Δ1290Δ1289)-<br>CA (ΔFEX) | NaF | ** | 0.0040 |
| *S. gordonii* | WT (3 species) | NaCl vs. NaF | ns | 0.8277 |
| | SM (Δ1290)-<br>CA (ΔFEX) | NaCl vs. NaF | ns | 0.7203 |
| | SM (Δ1289)-<br>CA (ΔFEX) | NaCl vs. NaF | ns | 0.6461 |
| | SM (Δ1290Δ1289)-<br>CA (ΔFEX) | NaCl vs. NaF | * | 0.0237 |
| | WT (3 species) vs.<br>SM (Δ1290)-<br>CA (ΔFEX) | NaF | ns | 0.8297 |
| | WT (3 species) vs.<br>SM (Δ1289)-<br>CA (ΔFEX) | NaF | ns | 0.2863 |
| | WT (3 species) vs.<br>SM (Δ1290Δ1289)- | NaF | ns | 0.2063 |
| | CA (ΔFEX) | | | |

**Supplementary Table 3:**
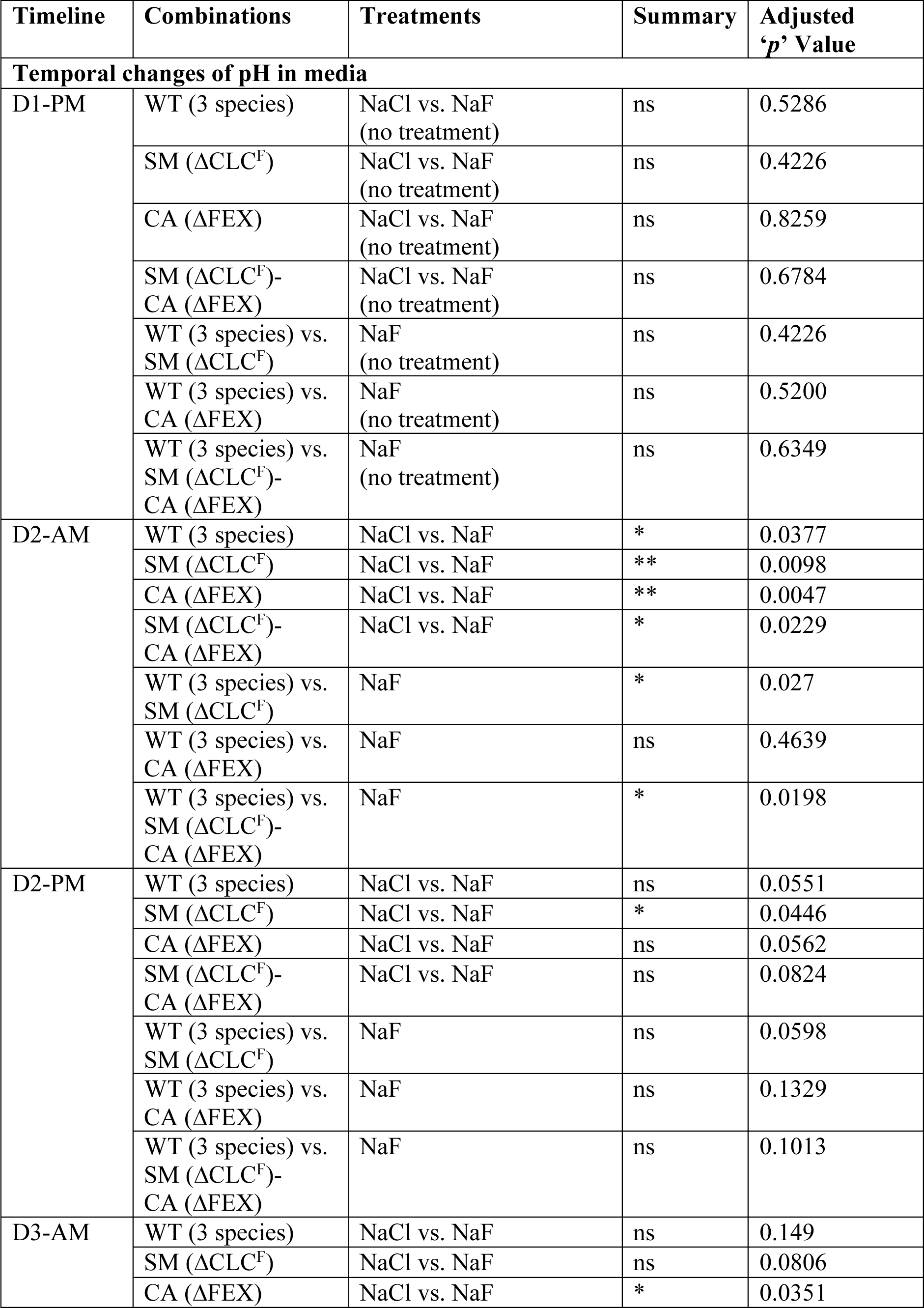
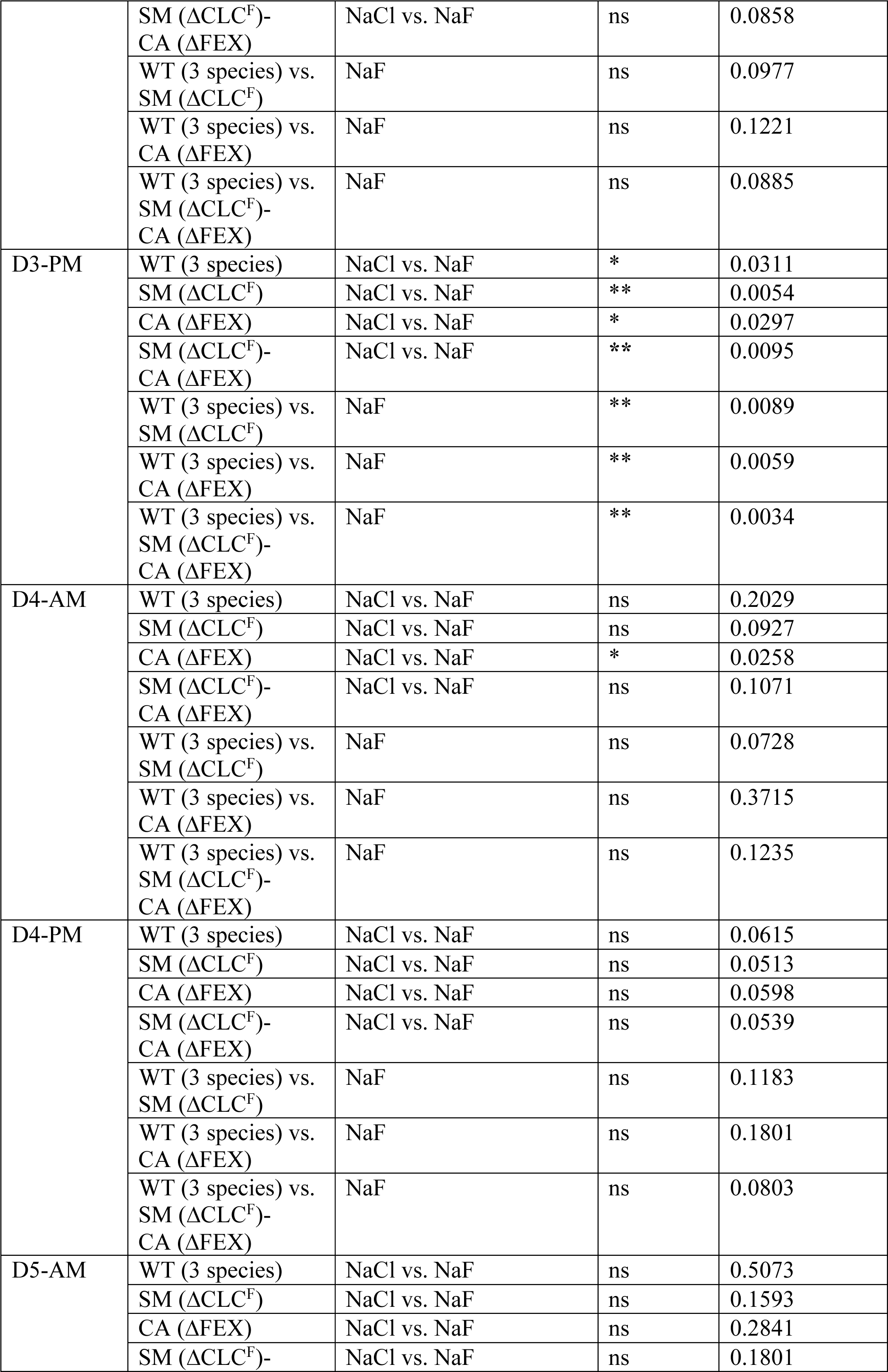
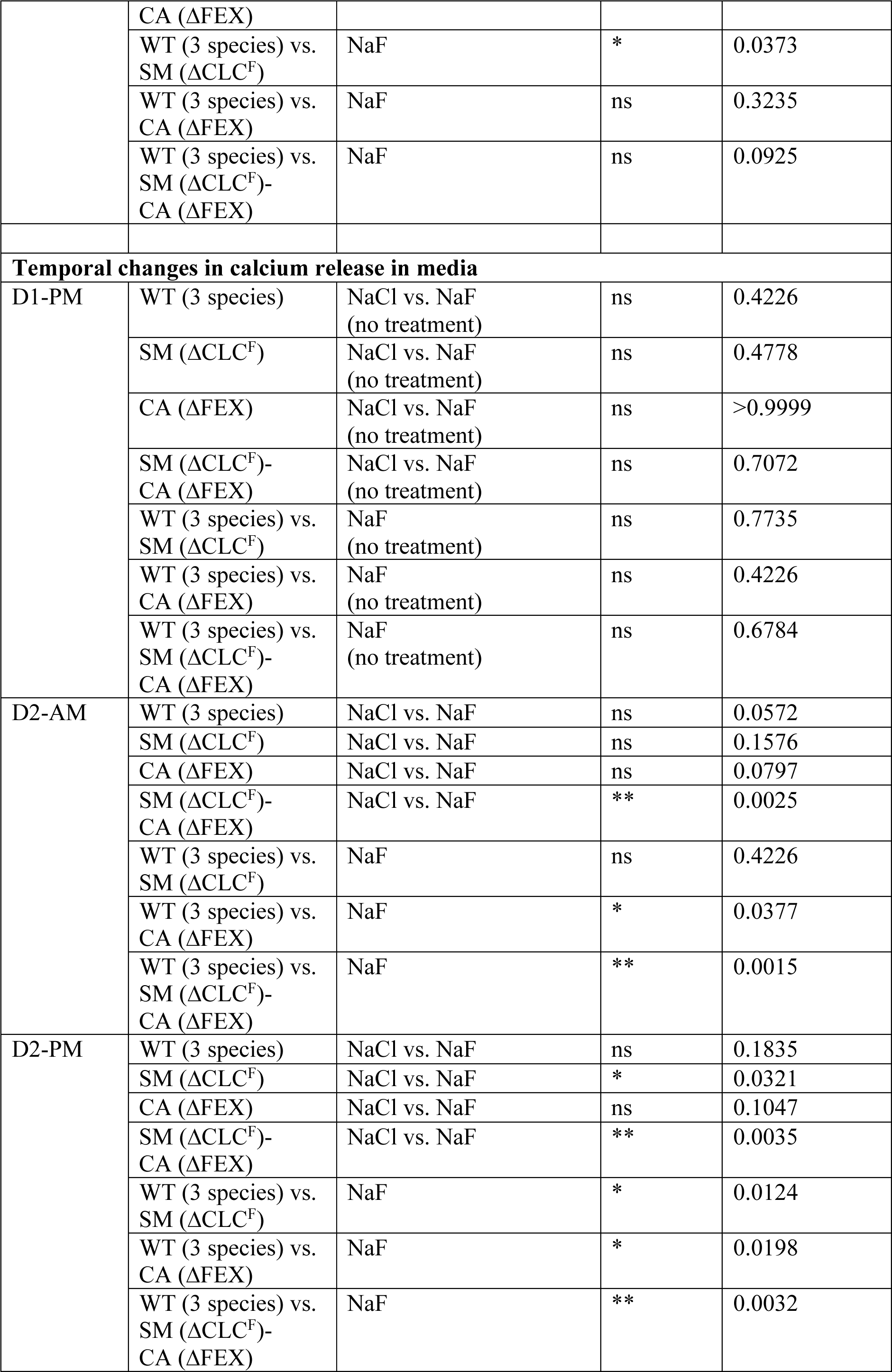
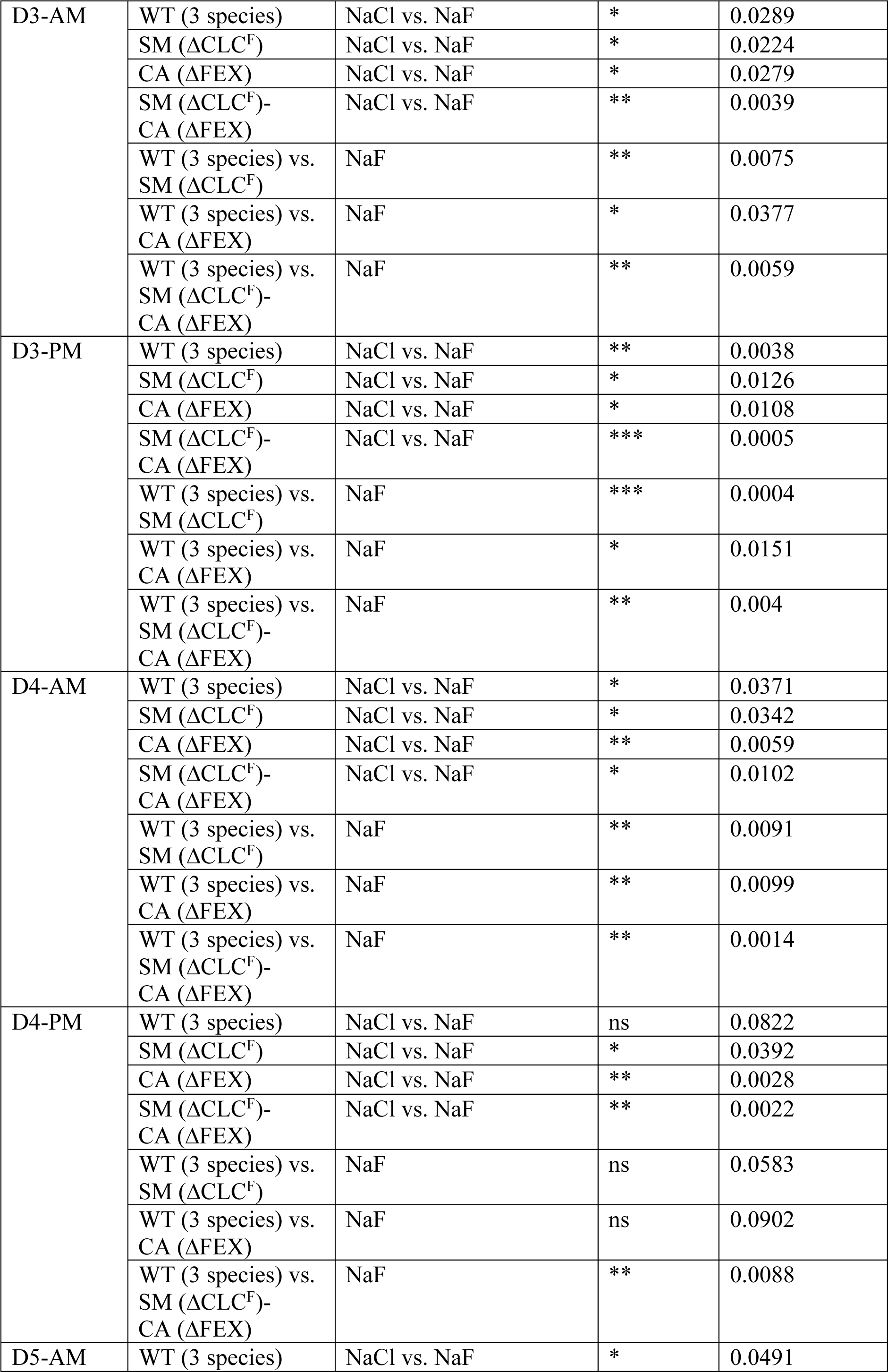
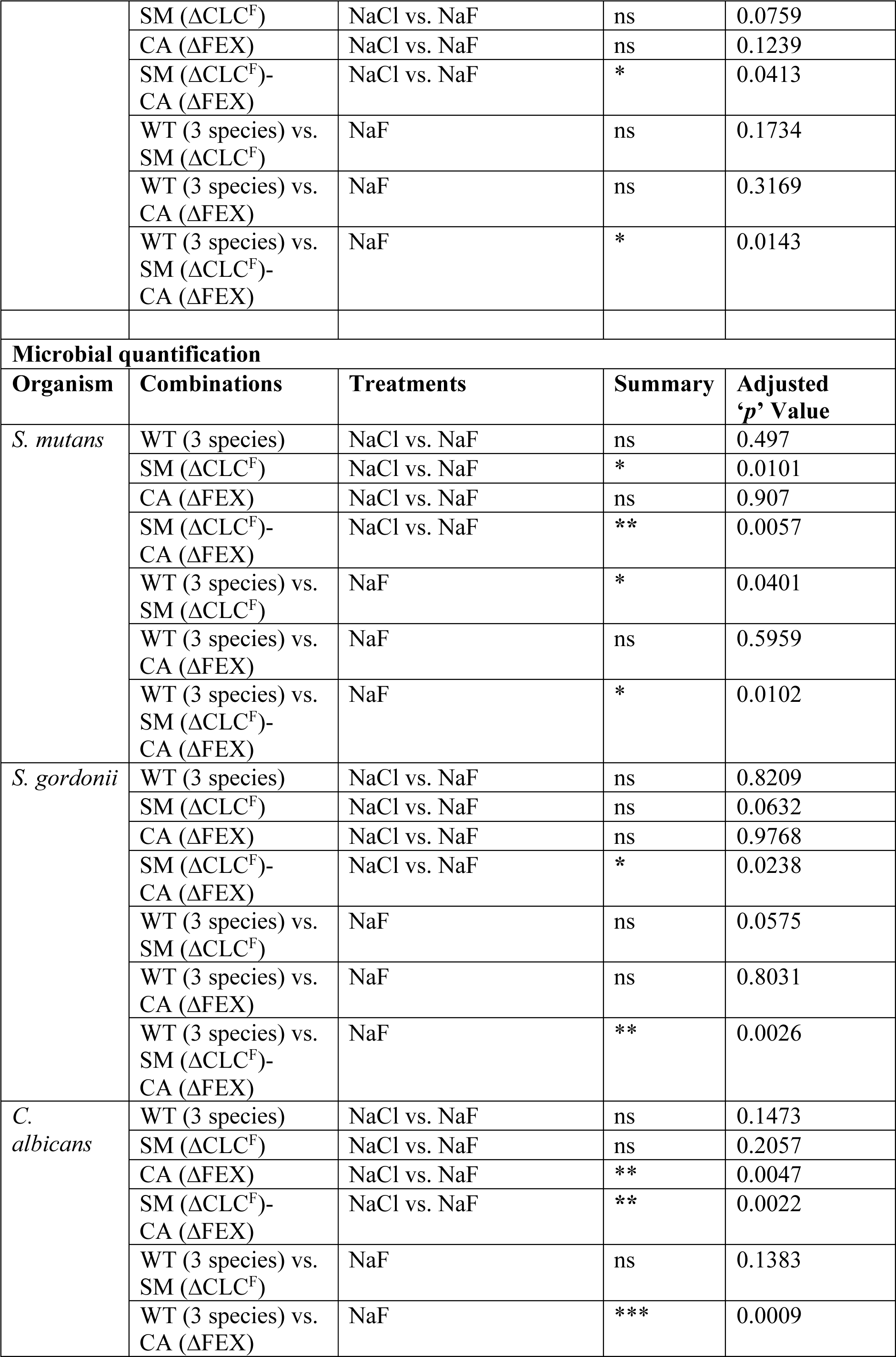
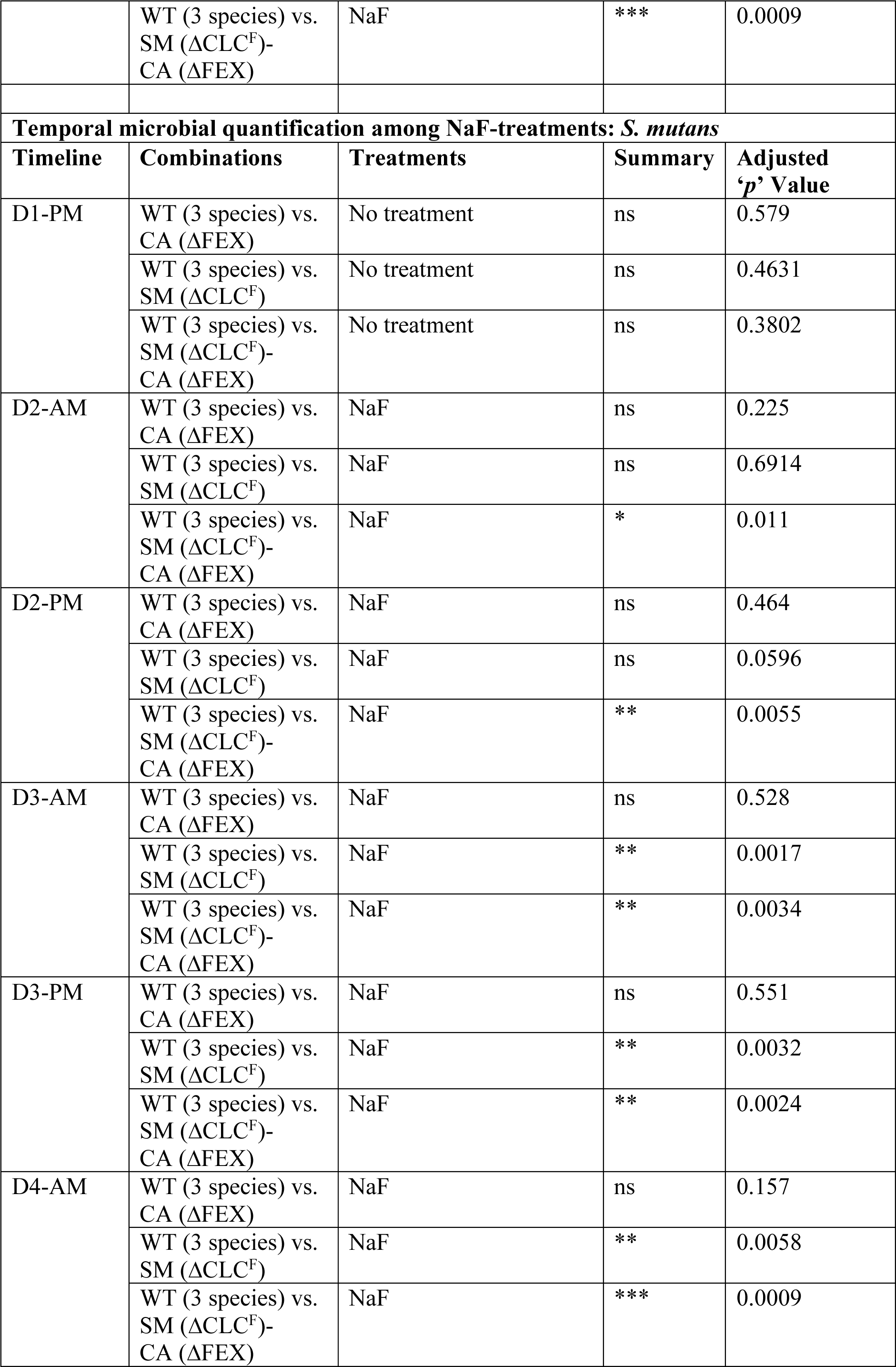
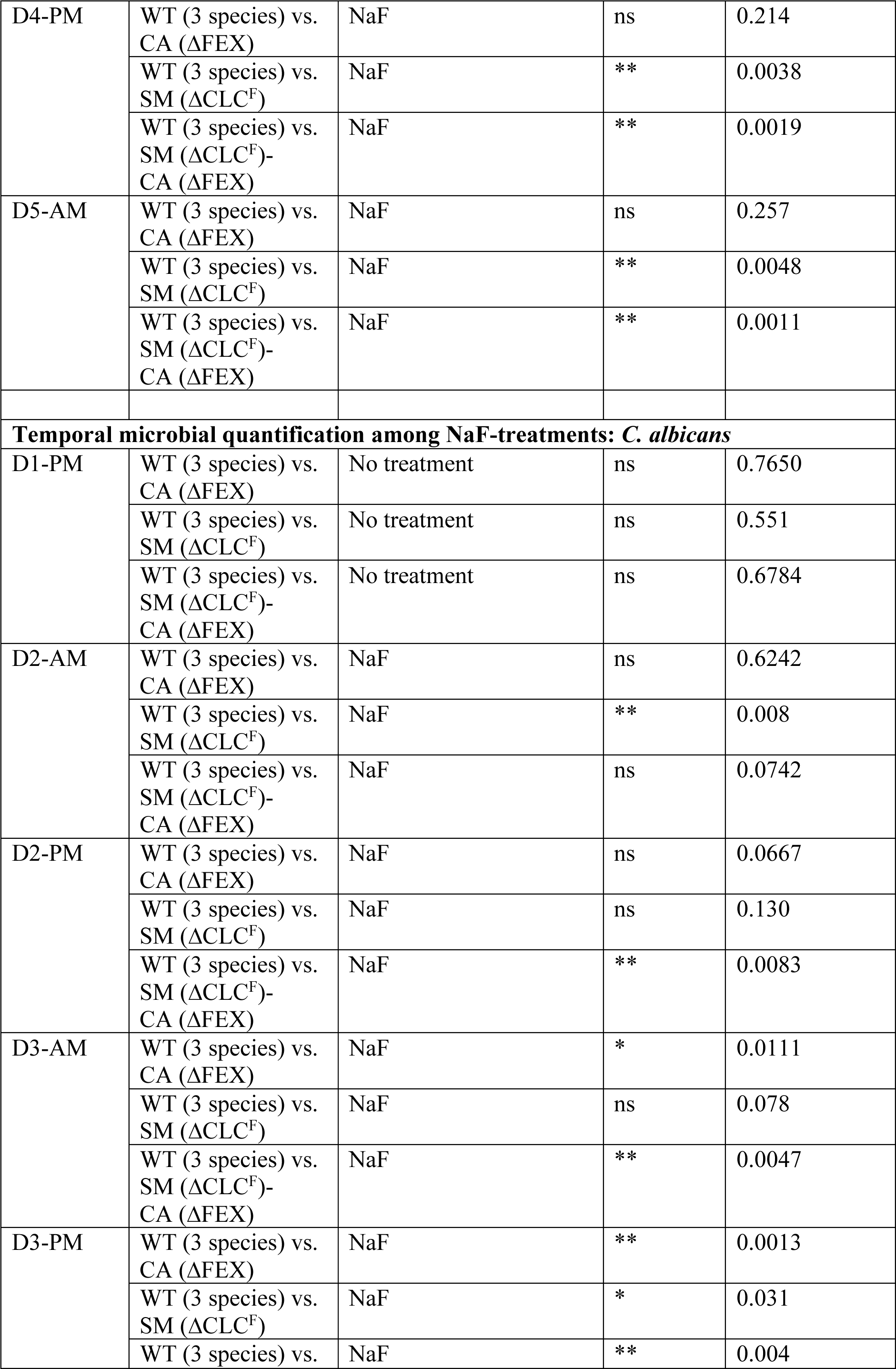
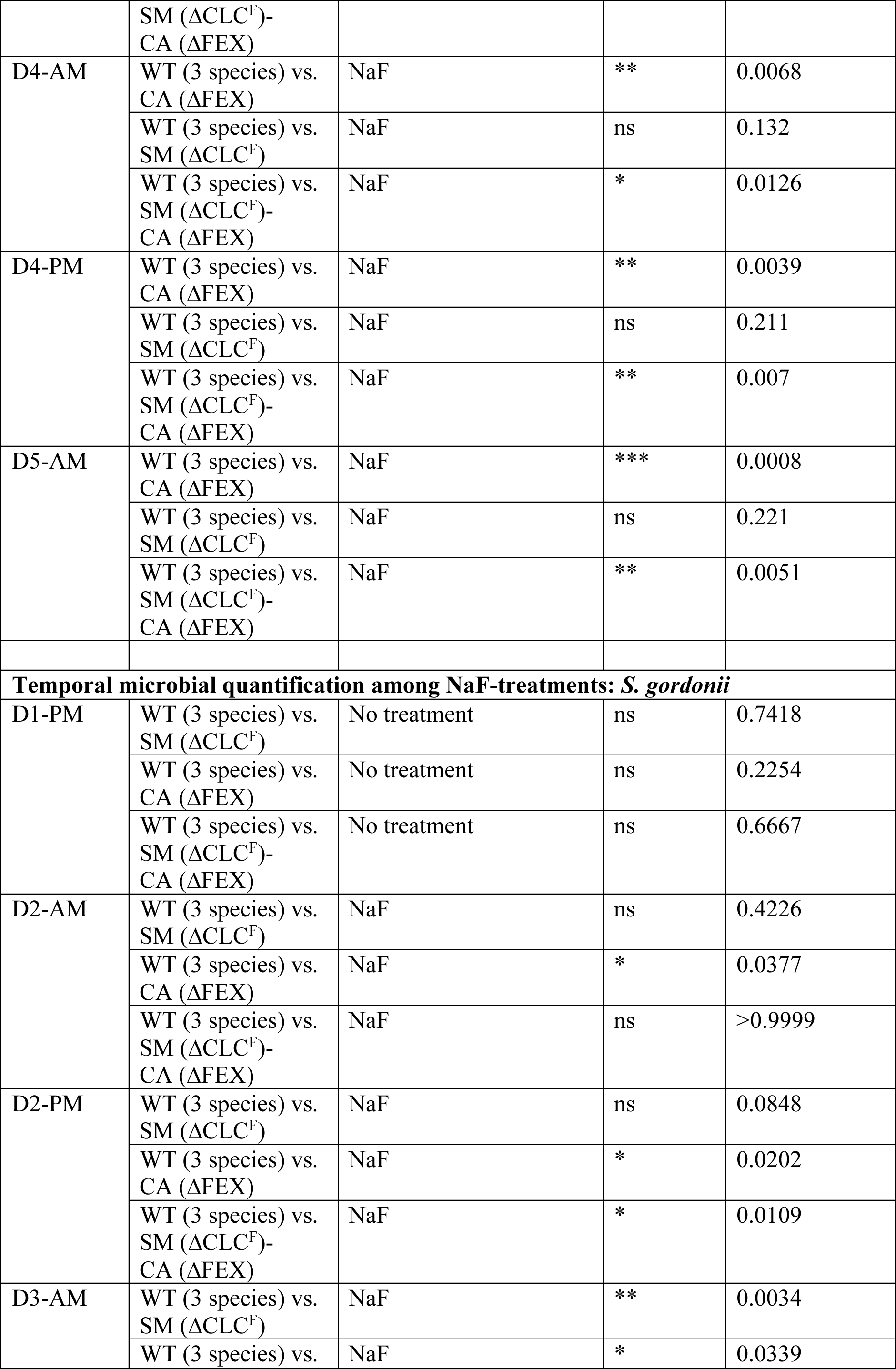
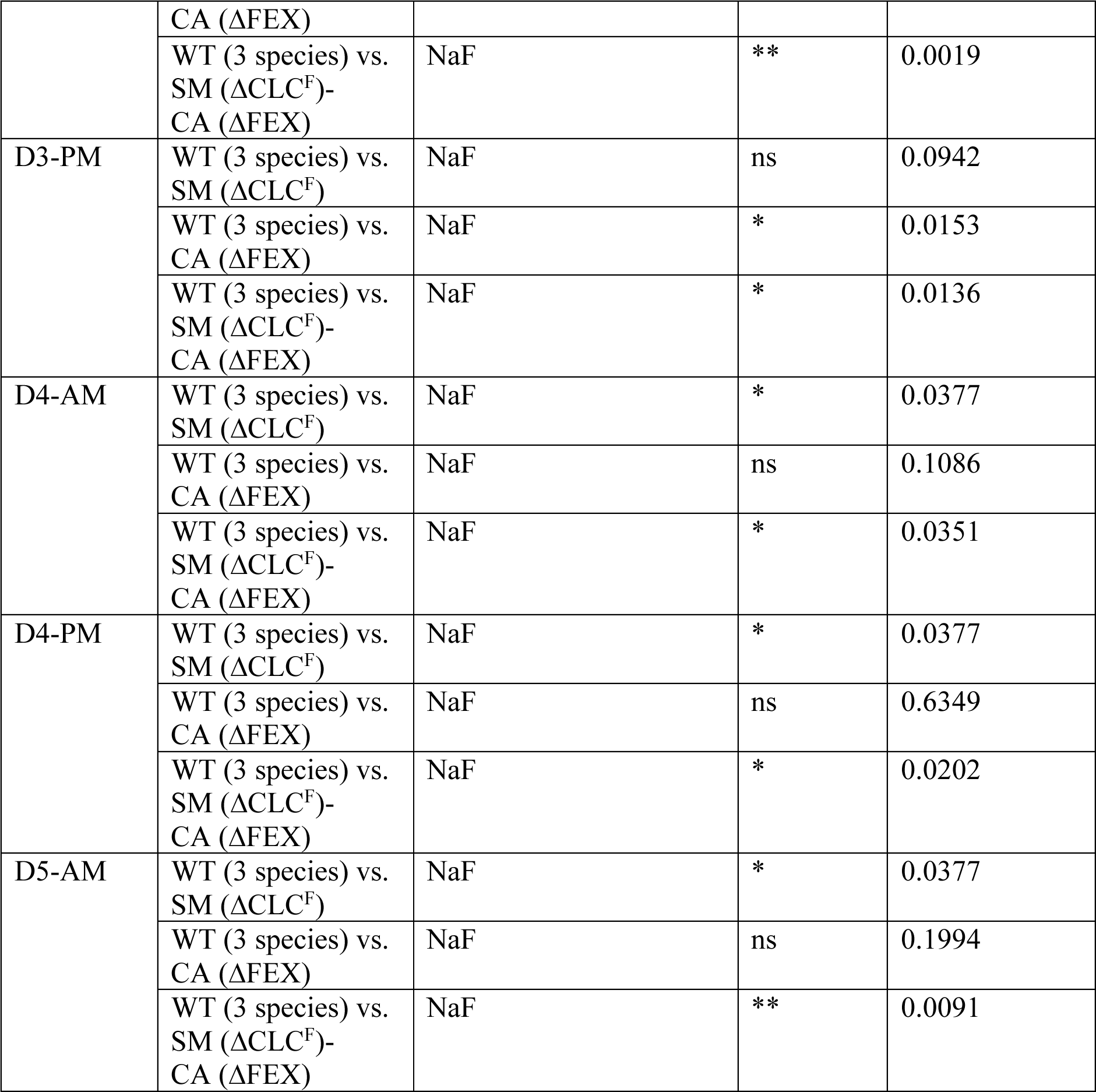
Statistical significance table for HA disc experiments. Significance was calculated using two-way analysis of variance (ANOVA) followed by Fisher’s LSD test implemented in GraphPad Prism 8. Statistical significance is represented as ‘*’ (*p*< 0.05), ‘**’ (*p*< 0.01) and ‘***’ (*p*< 0.001).

## Notes

### Competing Interest Statement

The authors have declared no competing interest.

